# A rapid and efficient learning rule for biological neural circuits

**DOI:** 10.1101/2021.03.10.434756

**Authors:** Eren Sezener, Agnieszka Grabska-Barwińska, Dimitar Kostadinov, Maxime Beau, Sanjukta Krishnagopal, David Budden, Marcus Hutter, Joel Veness, Matthew Botvinick, Claudia Clopath, Michael Häusser, Peter E. Latham

**Author notes:** Equal contribution (Eren Sezener, Agnieszka Grabska-Barwińska, Dimitar Kostadinov). Equal contribution (Claudia Clopath, Michael Häusser, Peter E. Latham).

## Abstract

The dominant view in neuroscience is that changes in synaptic weights underlie learning. It is unclear, however, how the brain is able to determine which synapses should change, and by how much. This uncertainty stands in sharp contrast to deep learning, where changes in weights are explicitly engineered to optimize performance. However, the main tool for that, backpropagation, has two problems. One is neuroscience related: it is not biologically plausible. The other is inherent: networks trained with this rule tend to forget old tasks when learning new ones. Here we introduce the Dendritic Gated Network (DGN), a variant of the Gated Linear Network, which offers a biologically plausible alternative to backpropagation. DGNs combine dendritic ‘gating’ (whereby interneurons target dendrites to shape neuronal responses) with local learning rules to yield provably efficient performance. They are significantly more data efficient than conventional artificial networks, and are highly resistant to forgetting. Consequently, they perform well on a variety of tasks, in some cases better than backpropagation. Importantly, DGNs have structural and functional similarities to the cerebellum, a link that we strengthen by using *in vivo* two-photon calcium imaging to show that single interneurons suppress activity in individual dendritic branches of Purkinje cells, a key feature of the model. Thus, DGNs leverage targeted dendritic inhibition and local learning – two features ubiquitous in the brain – to achieve fast and efficient learning.

## 1 Introduction

A hallmark of intelligent systems is their ability to learn. Humans, for instance, are capable of amazing feats – language acquisition and abstract reasoning being the most notable – and even fruit flies can learn simple reward associations [1,2]. It is widely believed that this learning is implemented via synaptic plasticity. But which synapses should change in response to, say the appearance of a reward, and by how much? This is especially hard to answer in humans, who have about 10^14^ synapses, but it is hard even in fruit flies, which have about 10^7^ – corresponding to 10 million adjustable parameters.

One answer to this question is known: introduce a loss function (a function that measures some aspect of performance, with higher performance corresponding to lower loss), compute the gradient of the loss with respect to the weights (find the direction in weight space that yields the largest improvement in performance), and change the weights in that direction. If the weight changes are not too large, this will, on average, reduce the loss, and so improve overall performance.

This approach has been amazingly successful in artificial neural networks, and has in fact driven the deep learning revolution [3]. However, the algorithm for computing the gradient in deep networks is not directly applicable to biological systems, as first pointed out by [4,5] (see also recent reviews [6–8]). There are several reasons for this. First, to implement backpropagation [9–11], referred to simply as backprop, neurons would need to know their outgoing weights. Second, backprop requires two stages: a forward pass (for computation) and a backward pass (for learning). Moreover, in the backward pass an error signal must propagate from higher to lower areas, layer by layer (Fig. 1A), and during that backward pass information from the forward pass must remain in the neurons. However, biological neurons do not know their outgoing weights, and there is no evidence for a complicated, time-separated backward pass.

**Figure 1:**
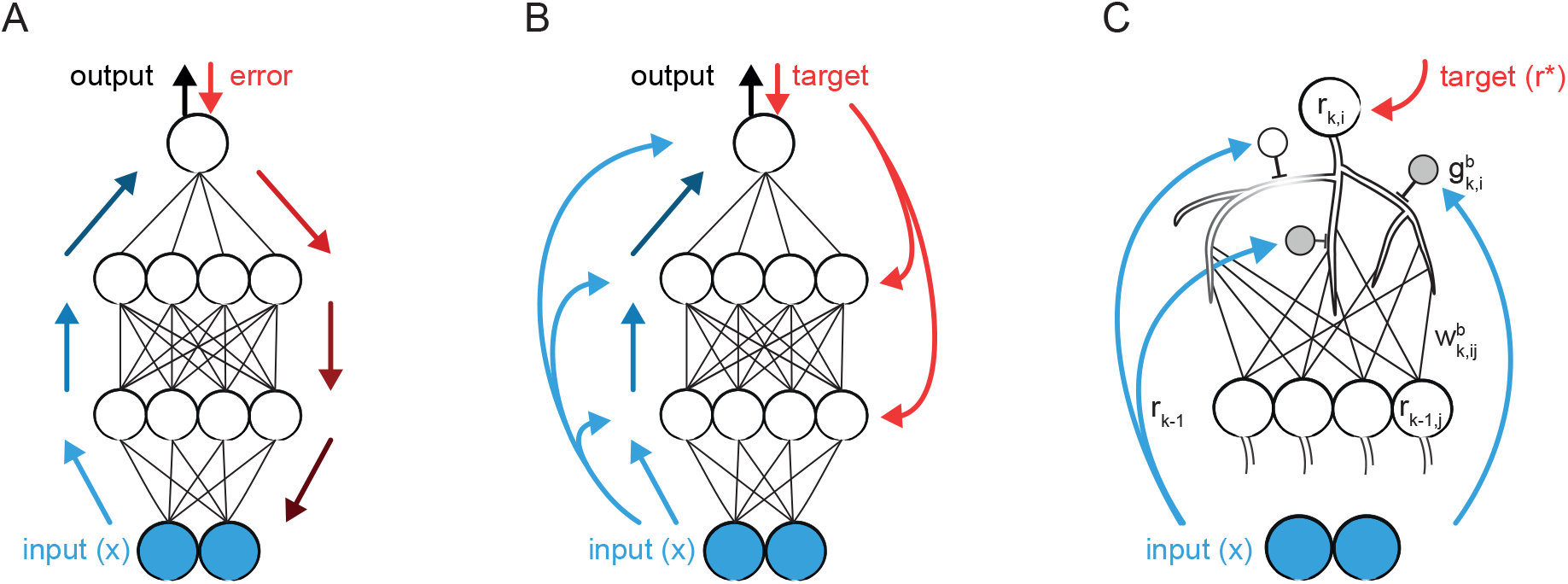
Comparison of multi-layer perceptrons (MLPs) and Dendrtic Gated Networks (DGNs). In all panels the blue filled circles at the bottom correspond to the input. A. MLP. Blue arrows show feedforward computations; red arrows show the error propagating back down. B. DGN. As with MLPs, information propagates up, as again shown by the blue arrows. However, rather than the error propagating down, each layer receives the target output, which it uses for learning. Connections from the input to each layer (light blue arrows) support the gating. C. A single postsynaptic neuron in layer *k* of a DGN, along with several presynaptic neurons in layer *k* – 1. Each branch gets input from all the presynaptic neurons (although this is not necessary), and those branches are gated on and off by inhibitory interneurons which receive external input. The white interneuron is active, so its corresponding branch is gated off, as indicated by the light gray branches; the gray neurons are not active, so their branches are gated on.

Backprop also leads to another problem, at least in standard deep learning setups: it adapts to the data it has seen most recently, so when learning a new task it forgets old ones [12]. This is known as catastrophic forgetting, and prevents networks trained with backprop to display the lifelong learning that comes so easily to essentially all organisms [13, 14].

Driven in part by the biological implausibility of backprop, there have been several proposals for architectures and learning rules that might be relevant to the brain. These include feedback alignment [15,16], creative use of dendrites [17,18], multiplexing [19], and methods in which the error signal is fed directly to each layer rather than propagating backwards from the output layer [20–28]. A particularly promising method that falls into the latter category is embodied in Gated Linear Networks [29, 30]. These networks, which were motivated from a machine learning rather than a neuroscience perspective, have obtained state-of-the-art results in regression and denoising [31], contextual bandit optimization [32], and transfer learning [33].

In Gated Linear Networks (GLNs), the goal of every neuron, irrespective of its layer, is to predict the target output based on the input from the layer directly below it. This is very different from backprop, in which neurons in intermediate layers extract features that make it easier for subsequent layers to predict the target (compare Figs. 1A and B). Gated Linear Networks are thus particularly suitable for biologically plausible learning: every neuron is essentially part of a shallow network, with no hidden layers, for which the delta rule [34] – a rule that depends only on presynaptic and postsynaptic activity – is sufficient to learn.

To implement these local learning rules, the target activity (a scalar) is sent to every neuron, in every layer of the network (Fig. 1B, red arrows). This is typical of a large class of learning rules [20,21,23–27]. Completely atypical, though, is the role of the external input. It is used for gating the weights: each neuron has a bank of weights at its disposal, and the external input determines which one from that bank is used. For example, a neuron might use one set of weights when the visual input contains motion cues predominantly to the right; another set of weights when it contains motion cues predominantly to the left; and yet another when there are no motion cues at all. (Note that this example is over-simplified: in practice the input is high dimensional, and the mapping from external input to the chosen set of weights contains very little structure; see Fig. 2C.)

**Figure 2:**
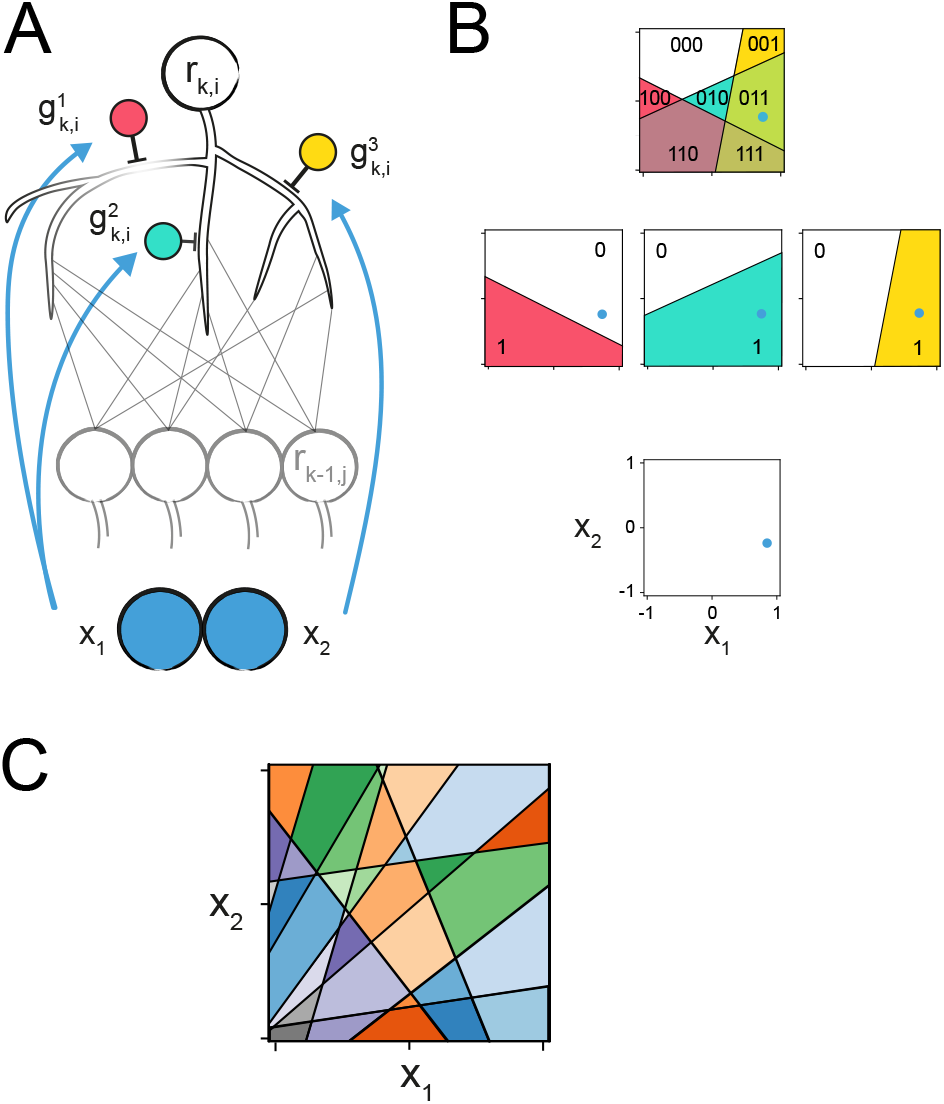
Random mapping (here implemented with so-called half-space gating, which we use throughout the paper; see Eq. (3)), shown for two dimensional input for clarity. (In realistic cases the input is, of course, high dimensional.) A. Two input neurons (blue) connect to three dendritic branches, and gate them either on or off. Each gate divides the two dimensional input into two half-spaces, one of which shuts down the gate, silencing the corresponding dendritic branch. B. For the input on this particular trial (blue dot in bottom square), the gate 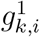 (red) is off, while 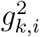 (teal) and 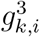 (yellow) are on, as indicated by the three squares with the corresponding colors. The top square shows a summary of the possible combinations of weights. Each of the seven regions has a different combination, making it possible to implement a large range of input-output mappings. C. A more realistic case of 10 branches. In DGNs, each coloured region corresponds to a linear combination of 10 sets of weights. This is in contrast to GLNs, which use a separate set of weights for each region.

Having a “look-up” table, in which each input corresponds to a particular set of weights, is inconsistent with what we see in the brain. However, we can attain the performance of Gated Linear networks by gating dendritic branches on and off, using inhibitory neurons, in an input-dependent manner (Figs 1C and 2). We thus replace the weight look-up mechanism of GLNs with linearly additive dendritic weights, and refer to these networks as Dendritic Gated Networks (DGNs). Perhaps surprisingly, the mapping from the input to the dendritic branches is completely random, so the input isn’t chosen to target specific branches (see, for example, Fig. 2C). This stands in sharp contrast to architectures where the gating is learned [35]. But the unlearned, random mapping is in fact a key ingredient, as it allows DGNs to represent essentially arbitrary nonlinear functions efficiently. Moreover, this gating makes DGNs especially resistant to forgetting. In particular, when data comes in separate “tasks”, DGNs can learn new ones without forgetting the old. Finally, the loss is a convex function of the weights for each unit (see Supplementary Information, “Convexity”), as it is in Gated Linear Networks [29]. Convexity is an extremely useful feature, as it enables DGNs, like the Gated Linear Networks on which they are based, to learn quickly.

Below we describe multi-layer Dendritic Gated Networks in detail – both the architecture and the learning rule. We then train them on four tasks: two on which vanilla feedforward networks trained with backprop typically exhibit catastrophic forgetting, and two relevant to the cerebellum. We map the proposed learning rule and the associated architecture to the cerebellum because 1) the climbing fibers provide a well-defined feedback signal; 2) its input-output function is relatively linear [36–38]; and 3) molecular layer interneurons could act as gates [39–48]. Finally, we demonstrate experimentally that a key prediction of the DGN – suppression of individual dendritic branches by interneurons – is observed in cerebellar Purkinje cell spiny branchlets *in vivo*. Thus, our theoretical and experimental results draw a specific link between learning in DGNs and the functional architecture of the cerebellum. The generality of the DGN architecture should also allow this algorithm to be implemented in a range of networks in the mammalian brain, including the neocortex.

## 2 Results

### 2.1 Dendritic Gated Networks

Dendritic Gated Networks, like conventional deep networks, are made up of multiple layers, with the input to each layer consisting of a linear combination of the activity in the previous layer. Unlike conventional deep networks, however, the weights are controlled by external input, via gating functions, denoted *g*(**x**); those functions are implemented via dendritic branches (Figs. 1B, C and 2). This results in the following network equations. The activity (i.e., the instantaneous firing rate) of the *i*^th^ neuron in layer *k*, denoted *r_k,i_*, is

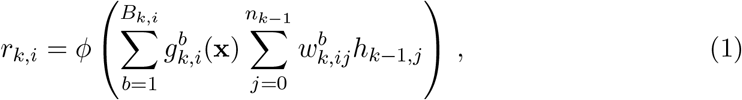

with the synaptic drive, *h*_*k*–1,*j*_, given in terms of *r*_*k*–1,*j*_ as

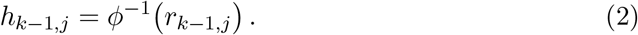

Here *ϕ*(·) is the activation function (either identity or sigmoid), *r_k,i_* is the activity of *i*^th^ neuron in layer *k* (with *r*_*k*,0_ set to 1 to allow for a bias), *B_k,i_* is the number of branches of neuron *i* in layer *k*, 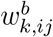 is the weight from neuron *j* in layer *k* – 1 to the *b*^th^ branch of neuron *i* in layer *k*, *n_k_* is the number of neurons in layer *k*, and 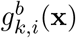 is the binary gating variable; depending on the external input, **x** (taken to be an *n*-dimensional vector), it’s either 1 (in which case the *b*^th^ branch of the *i*^th^ neuron is gated on) or 0 (in which case it’s gated off). There are *K* layers, so *k* runs from 1 to *K*. The input to the bottom layer is **x** – the same as the input to the gating variable.

The mapping from the input, **x**, to the gating variable, 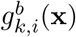, is not learned; instead, it is pre-specified, and does not change with time. In all of our simulations we use random half-space gating [29]; that is,

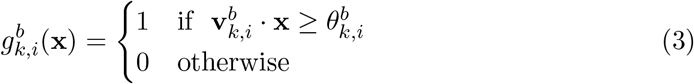

where 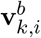 and 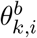 are sampled randomly and kept fixed throughout learning (see Methods), and “·” is the standard dot product.

Note that a Dendritic Gated Network with one branch reduces to a Gated Linear Network. With a caveat: in the original formulation [29], Gated Linear Networks had a nonzero weight for each input, **x**, which is not the case if weights are completely gated off for one of the half spaces (because in that case the weights are zero). For a detailed description of the difference between GLNs and DGNs, see Supplementary Information, “Difference between GLNs and DGNs”.

In Dendritic Gated Networks, the goal of each neuron is to predict the target output, denoted *r*\* (which is a function of **x**; we suppress the **x**-dependence to reduce clutter). To do that, the weights, 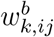, are modified to reduce the loss, *ℓ_k_*(*r*\*, *r_k,i_*). For weight updates we use gradient descent,

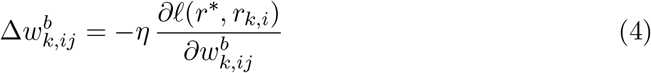

where *η* is the learning rate, and the updates are performed after each sample. The form of the loss can influence both the speed of learning and the asymptotic performance, but conceptually we should just think of it as some distance between *r*\* and *r_k,i_*. In the simplest case, which is suitable for regression, *ϕ* is the identity (*r_k,i_* = *h_k,i_*) and the loss is quadratic,

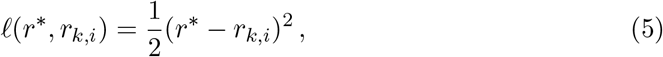

so the update rule is

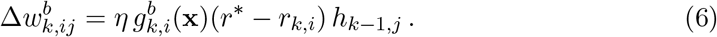

This has the form of a gated version of the delta rule [34]. For classification, a different loss function is more appropriate. However, the update rule still has the form of a gated version of the delta rule; see Methods, Sec. 4.1, for details.

### 2.2 Simulations

Equations (1) and (3) for the network dynamics and Eq. (4) for learning constitute a complete description of our model. For a given problem, we just need to choose a target input-output relationship (a mapping from **x** to *r*\*) and specify the loss functions, *ℓ*(*r*\*, *r_k,i_*). Here we consider four tasks. The first two, designed to illustrate the resistance of DGNs to catastrophic forgetting, are classification tasks, for which we use a sigmoid activation and cross-entropy loss (Methods, Sec. 4.1); the second two, which are relevant to the cerebellum, are regression tasks, for which we use an identity activation and quadratic loss, as just described.

#### DGNs can mitigate catastrophic forgetting

Animals are able to acquire new skills throughout life, seemingly without compromising their ability to solve previously learned tasks [13, 14]. Standard networks do not share this ability: when trained on two tasks in a row, they tend to forget the first one. This phenomenon, known as “catastrophic forgetting”, is an old problem [49–51], and many algorithms have been developed to address it. These typically fall into two categories. The first involves replaying tasks previously seen during training [51–53]. The second involves explicitly maintaining additional sets of model parameters related to previously learned tasks. Examples include freezing a subset of weights [54, 55], dynamically adjusting learning rates [56], and augmenting the loss with regularization terms with respect to past parameters [57–59]. A limitation of these approaches (aside from additional algorithmic and computational complexity) is that they require task boundaries to be provided or accurately inferred.

Unlike contemporary neural networks, the DGN architecture and learning rule is naturally robust to catastrophic forgetting without any modifications or knowledge of task boundaries (something that has been shown for Gated Linear Networks as well [30]). In Fig. 3 we illustrate, on a simple task, the mechanism behind this robustness, and show how it differs from a standard multi-layer perceptron; details are given in the caption.

**Figure 3:**
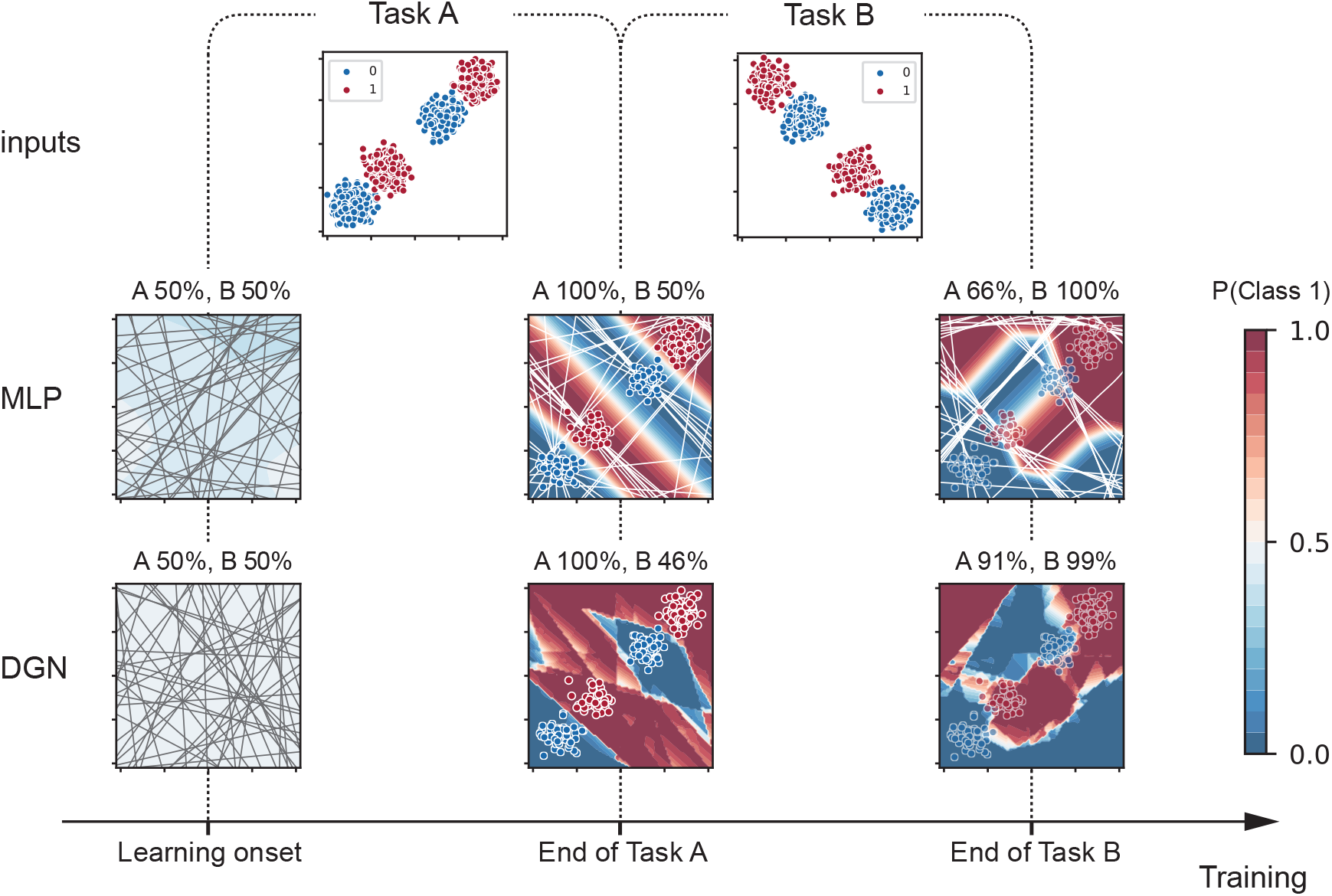
Comparison of the DGN to a standard multi-layer perceptron (MLP) trained with backprop. Each point on the square has to be classified as “blue” (class 0) or “red” (class 1). We consider a scenario common in the real world (but difficult for standard networks): the data comes in two separate tasks, as shown in the first row. We trained a 2-layer MLP (second row) and a 2-layer DGN (third row) on the two tasks. The output of the network is the probability of each class, as indicated by color; the percentages report the accuracy for each of the tasks. The MLP uses ReLU activation functions, so each neuron has an effective gating; the boundaries of those gates are shown in gray. The boundaries move with learning, and are plotted at the end of training of each of the tasks (white lines). The boundaries of the DGN do not move, so we plot them only in the first column. After training on Task A, most of the boundaries in the MLP are aligned at −45 degrees, parallel to the decision boundaries, which allows the network to perfectly separate the two classes. In the DGN, the boundaries do not change, but the network also perfectly separates the two classes. However, after training on Task B, the DGN retains high performance on Task A (91%), while the MLP’s performance drops to 66%. That’s because many of the boundaries changed to the orthogonal direction (45 degrees). For the DGN, on the other hand, changes to the network were much more local, allowing it to retain the memory of the old task (see samples from Task A overlaid on all panels) while accommodating the new one. The MLP has 50 neurons in the hidden layer; the DGN has 5 neurons, each with 10 dendritic branches, in the hidden layer.

To demonstrate robustness to catastrophic forgetting on a more challenging task, we train a DGN on the pixel-permuted MNIST continual learning benchmark [57, 60]. In this benchmark, the network has to learn random permutations of the input pixels, with the random permutation changing every 60,000 trials (see Methods for additional details). We compare the DGN to a multi layer perceptron (MLP) with and without elastic weight consolidation (EWC) [57], the latter a highly-effective method explicitly designed to prevent catastrophic forgetting by storing parameters of previously seen tasks. Although elastic weight consolidation is effective, it requires a very complicated architecture. In addition, it must be supplied with task boundaries, so it receives more information than the DGN.

Because MNIST has 10 digits, we train 10 different DGNs. This could be reduced to 4 networks (in general log_2_ of the number of outputs) by using a more efficient code – one in which each network divides the 10 digits into two classes. Alternatively, we could use a single DGN where each unit has a 10 dimensional output corresponding to the class probabilities. However, this is not biologically plausible, so we did not use it. Each of the 10 networks contains 3 layers, with 100, 20, and 1 neuron per layer, and there are 10 dendritic branches per neuron. The targets are categorical (1 if the digit is present, 0 if it is not), so we use binary cross-entropy rather than quadratic loss (see Methods, Sec. 4.1). We use 1000, 200, and 10 neurons per layer for the MLP (so that the number of weights match, approximately, the number weights in the DGN), with categorical cross entropy loss, both with and without elastic weight consolidation, and optimize the learning rates separately for each network.

Figure 4 shows the learning and retention performance of the DGN, with the MLP and EWC networks included primarily as benchmarks (neither is biologically plausible). In Fig. 4A we plot performance on each task for the three networks; as can be seen, performance is virtually identical. In Fig. 4B we investigate resistance to forgetting, by plotting the performance on the first task as the nine subsequent tasks are learned. The EWC network retains its original performance almost perfectly, the MLP forgets rapidly, and the DGN is in-between. It is not surprising that the EWC does well, as it was tailored to this task, and in particular it was explicitly given task boundaries. Somewhat more surprising is the performance of the DGN, which had none of these advantages but still forgets much more slowly than the MLP. The DGN also learns new tasks more rapidly than either the EWC or MLP networks (Supplementary Figure S3), because the loss is convex and learning is local.

**Figure 4:**
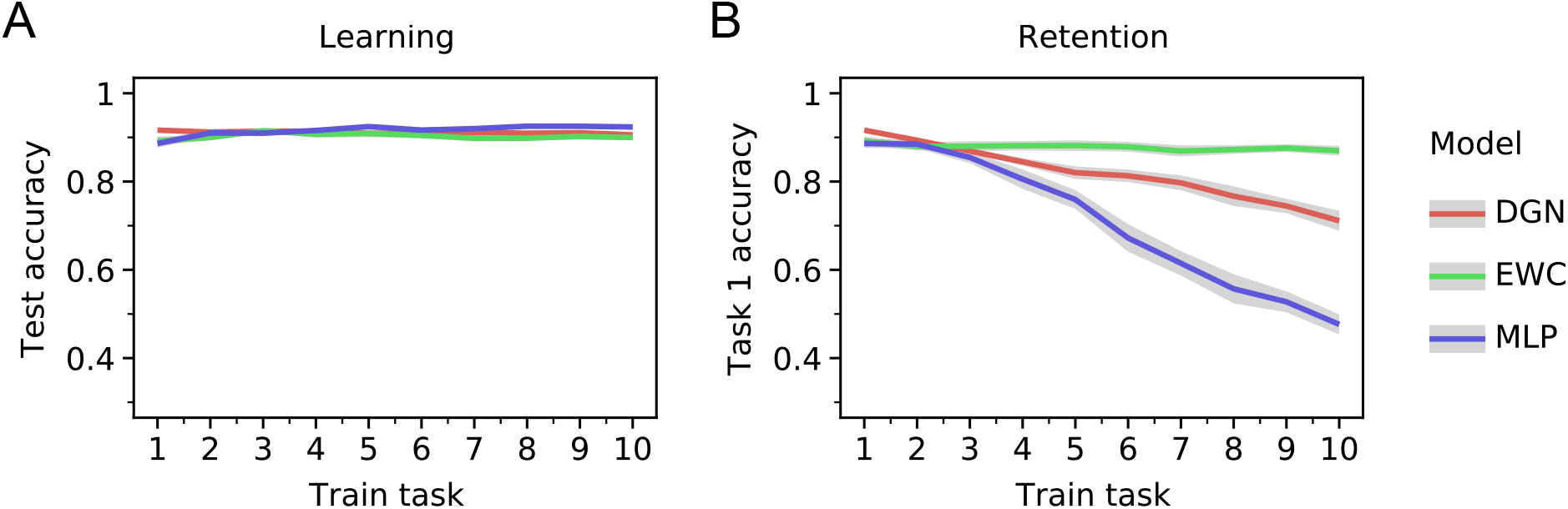
Learning and retention on the permuted MNIST task. The tasks are learned sequentially in a continual learning setup. A. Performance (on test data) for each of the 10 tasks, where a “task” corresponds to a random permutation of the pixels. B. Performance on the first task after each of nine new tasks is learned. As discussed in the main text, the MLP is especially bad at this task. The EWC is much better, to a large extent because it received extra information: the task boundaries. Even though the DGN was not given that information, it forgets a factor of two more slowly than the MLP. Error bars in both plots denote 95% confidence intervals over 20 random seeds.

#### Mapping DGNs to the Cerebellum

For the next two simulations we consider computations that can be mapped onto cerebellar circuitry. We focus on the cerebellum for several reasons: it is highly experimentally accessible; its architecture is well characterized; there is a clear feedback signal to the Purkinje cells (the cerebellar neurons principally involved in learning); its input-output function is relatively linear [36–38]; and molecular layer interneurons play a major role in shaping Purkinje cell responses [39–45,47], and can influence climbing fiber-mediated dendritic calcium signals in Purkinje cells [46, 48, 61].

Both classic and more modern theoretical studies in the cerebellum have focused on the cerebellar cortex, modelling it as a one-layer feedforward network [62–66]. In this view, the parallel fibers project to Purkinje cells, and their synaptic weights are adjusted under the feedback signal from the climbing fibers. This picture, however, is an over-simplification, as Purkinje cells do not directly influence downstream structures. Instead, they project to the cerebellar nucleus, which constitutes the output of the cerebellum (see Fig. 5). The fact that Purkinje cells form a hidden layer, combined with the observed plasticity in the Purkinje cell to cerebellar nucleus synapses [67–71], means most learning rules tailored to one-layer networks, including the delta rule, cannot be used to train the network.

**Figure 5:**
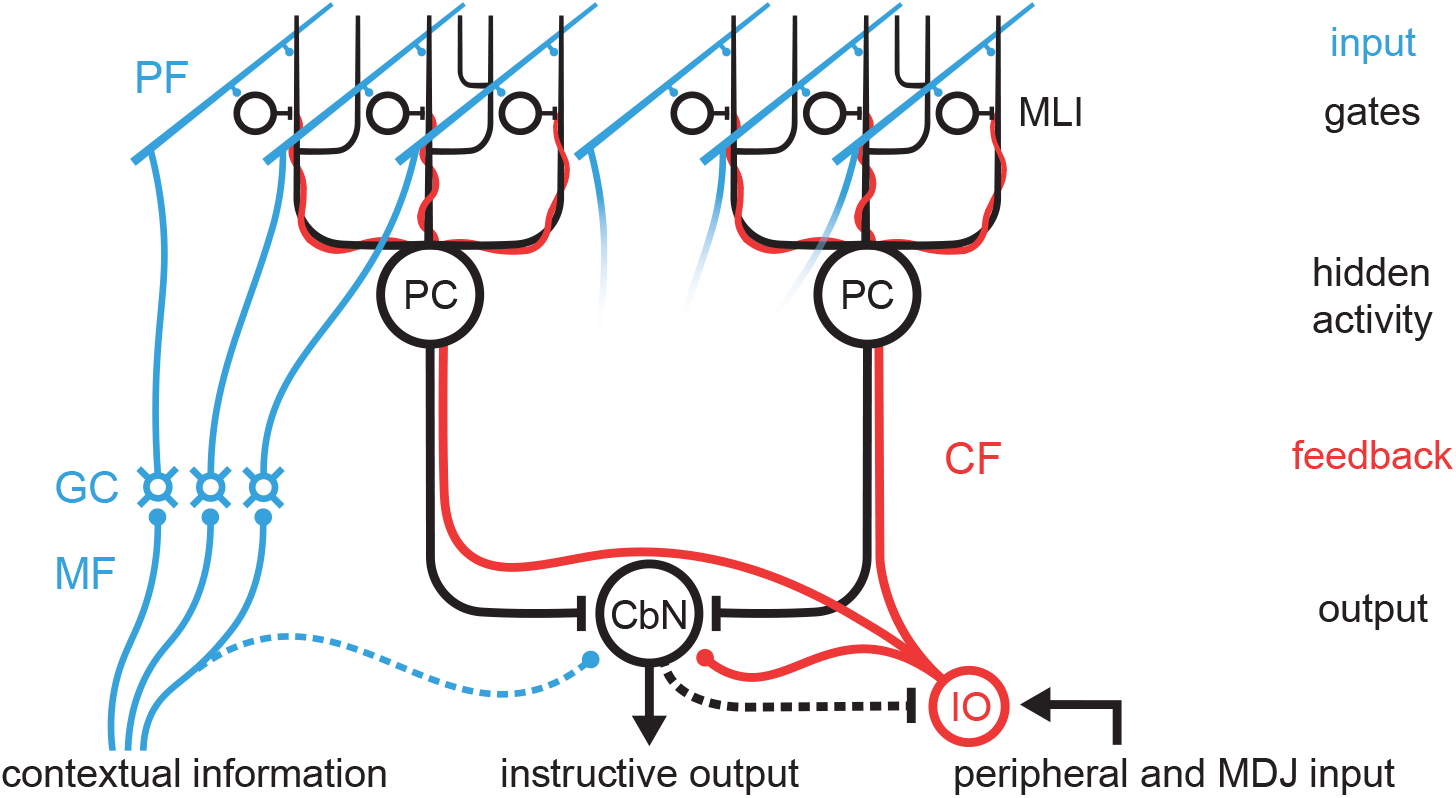
The cerebellum as a two layer DGN. Contextual information from the mossy fiber/granule cell (MF/GC) pathway is conveyed as input to the network via parallel fibers (PFs) that form synapses onto both the dendritic branches of Purkinje cells and molecular layer interneurons (MLIs). The inhibitory MLIs act as input-dependent gates of Purkinje cell dendritic branches. Purkinje cells converge onto the cerebellar nuclear neurons (CbNs), which constitute the output of the cerebellar network. The climbing fibers (CFs, red) originating in the inferior olive (IO) convey the feedback signal that is used to tune both the Purkinje cells, based on which inputs are gated on or off, and also the CbNs. Excitatory and inhibitory connections are depicted as round- and T-ends, respectively. Dashed lines represent connections not included in the model.

We propose instead that the cerebellum acts as a two layer DGN comprised of Purkinje cells as the first, hidden layer and the cerebellar nucleus as the second, output layer (Fig. 5). Parallel fibers provide the input to both the input layer (Purkinje cells) and the gates, represented by molecular layer interneurons, that control learning in individual Purkinje cell dendrites. For the output layer of the DGN (which consists of one neuron), we use a non-gated rather than a gated neuron, as the unique biophysical features of cerebellar nuclear neurons allow them to integrate inputs approximately linearly [72]. The climbing fibers provide the feedback signal to Purkinje cells and cerebellar nuclear neurons. In our formulation, climbing fiber feedback signals the target, allowing each neuron to compute its own local error by comparing the target to its output (*r_k,i_*). This formulation is a departure from the strict error-coding role that is traditionally attributed to climbing fibers, but is consistent with a growing body of evidence that climbing fibers signal a variety of sensorimotor and cognitive predictions [73].

#### DGNs can learn inverse kinematics

The cerebellum is thought to implement inverse motor control [74, 75]. We therefore applied our proposed DGN network to the SARCOS benchmark [76], which is an inverse kinematics dataset from a 7 degree-of-freedom robot arm (Fig. 6). The goal is to learn an inverse model, and predict 7 torques given the joint positions, velocities, and accelerations for each of the 7 joints (corresponding to a 21 dimensional input).

**Figure 6:**
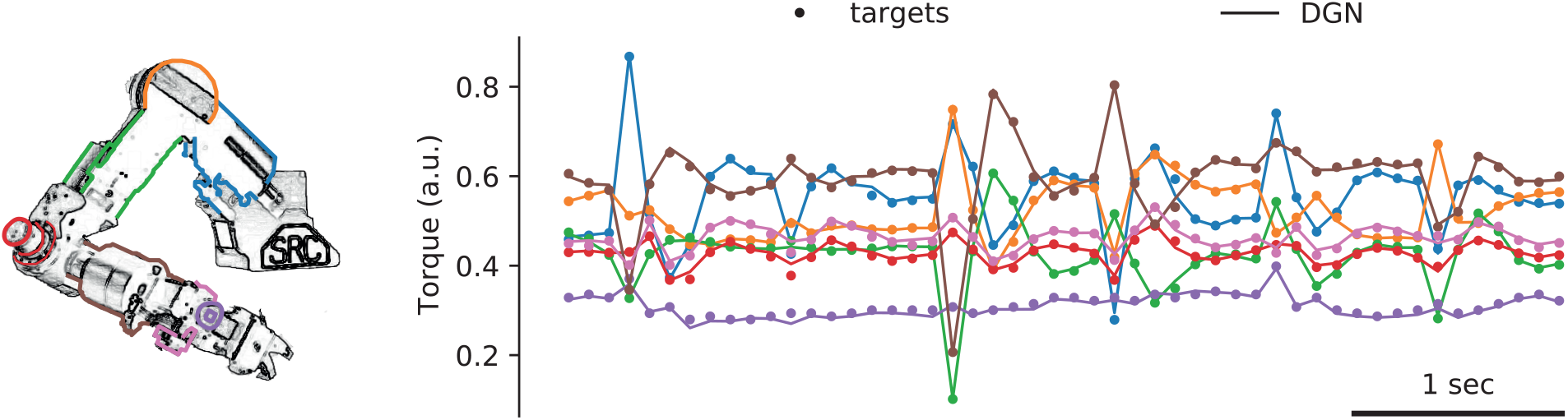
Sarcos experiment. DGNs can solve a challenging motor control task: predicting torques from the proprioceptive input. The data comes from a SARCOS dexterous robotic arm [76], pictured on the left. The inputs are position, velocity and acceleration of the 7 joints (a 21 dimensional variable); the targets are the desired torques (7 dimensional). Example targets (normalized to keep the training data between 0 and 1) are shown with dots, the lines are the output of our network. Performance is very good; only rarely is there a visible difference between the dots and the lines.

The target output, *r*\*, is the desired torque, given the 21-dimensional input. There are seven joints, so we train seven different networks, each with its own target output. We use DGN networks with 20 Purkinje cells, and minimize a quadratic loss (5). Since this is a relatively hard task, performance depends strongly on the number of branches. In Fig. 6 we plot the target torques for each joint (dots) along with the predictions of the DGN (lines; chosen for ease of comparison as there is no data between the points) for 500 branches. The lines follow the points reasonably closely, even when there are large fluctuations, indicating that the DGN is faithfully predicting torques. The performance of our network (mean squared error on test data in the original torque units) is comparable to that of most existing machine learning algorithms (Supplementary Table S1) while using fewer samples to learn. In Supplementary Fig. S2 we show the equivalent plot for 5, 50 and 5000 branches. Even at 5 branches performance is reasonable, while at 5000 we exceed the performance of almost all existing machine learning algorithms.

#### Vestibulo-ocular reflex, and adaptation to gain changes

To maintain a stable image on the retina during head movements, when an animal moves its head it moves its eyes in the opposite direction. This is known as the vestibulo-ocular reflex (VOR), and a key feature of it is that it’s plastic: animals can adapt quickly when the relationship between the head movement and visual feedback is changed, as occurs as animals grow or are given corrective lenses. VOR gain adaptation relies critically on the cerebellum, and has been used to study cerebellar motor learning for decades [77–81]. This is an easy task to learn – almost any network, including a linear one, can achieve high performance on it. We consider it primarily because it is a very common cerebellar task.

We applied our DGN network to the VOR in a regime where the gain occasionally changes abruptly. The gain, denoted *G*, is the ratio of the desired eye velocity to the head velocity (multiplied by −1 because the eyes and head move in opposite direction, to keep with the convention that the gain is reported as a positive number). When the gain is (artificially) changed, at first animals move their eyes at the wrong speed, but after about 15 minutes they learn to compensate [79,80].

We trained our network on a head velocity signal of the form

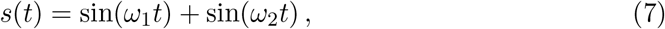

with *ω*_1_ = 13.333 and *ω*_2_ = 20.733 (corresponding to 2.12 and 3.30 Hz, respectively). This signal was chosen to mimic, approximately, the irregular head velocities encountered in natural viewing conditions. Following Clopath et. al. [82], we assumed that the Purkinje cells receive delayed versions of this signal. The *i*^th^ input signal, *x_i_*(*t*), which arrives via the parallel fibers, is modelled as

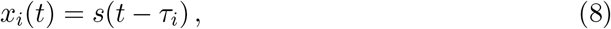

with delays, *τ_i_*, spanning the range 50-300 ms. The cerebellum needs to compute the scaled version of the eye velocity: *r*\*(*t*) = *Gs*(*t*) (as mentioned above, the actual eye movement is –*r*\*(*t*), but we follow the standard convention). Learning was online, and we updated the weights every 500 ms, to approximately match the climbing fiber firing rate [83].

The DGN contained 20 Purkinje cells, with 10 branches each; these project to one output neuron (corresponding to the cerebellar nucleus), which was linear and not gated. As a baseline, we trained an MLP with the same number of weights (resulting in 200 hidden neurons). We used quadratic loss for both the DGN and the MLP and, as in [82], we assumed *n* = 100 parallel fibers and a single output. Each branch received input from all 100 parallel fibers. Gating (Eq. (3)) was controlled by *x_i_*(*t*) (given in Eq. (8)), reflecting the parallel fiber influence on molecular layer interneurons (Fig. 5); see Methods for details. Given the timescale of the signal (2-3 Hz), any individual branch was gated on for about 500 ms at a time. The networks were pre-trained on a gain, *G*, of 1. We implemented four jump changes: first to 0.7, then back to 1.0, then to 1.3, and, finally, back to 1.0; in all cases, for 30 minutes at each gain (Fig. 7A).

**Figure 7:**
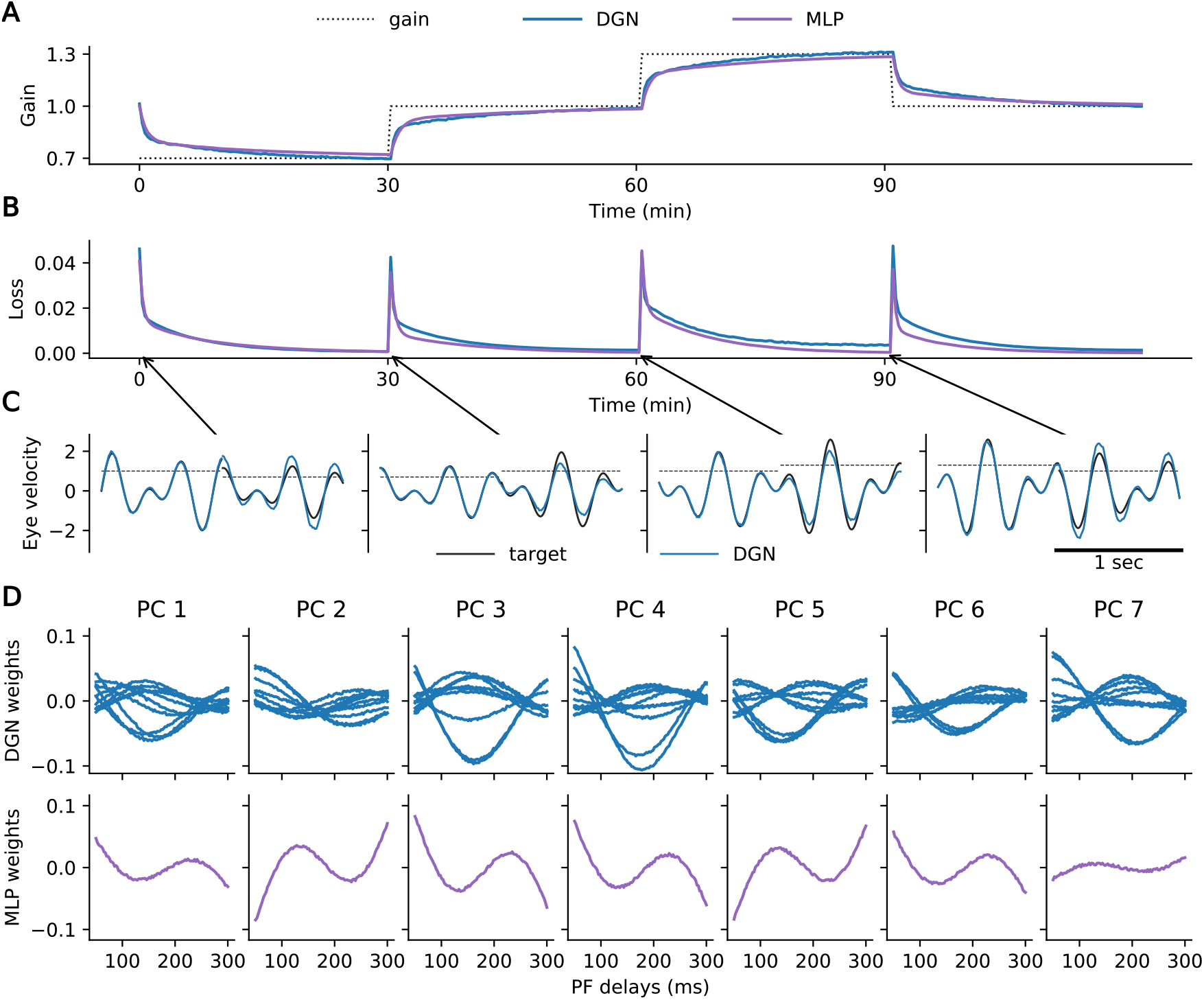
VOR adaptation task. We trained the networks on gain *G* = 1, then changed the gain every 30 minutes. Results are shown for the Dendritic Gated Network (DGN) and a multi-layer perceptron (MLP). A. Dashed lines are true gain versus time; blue and purple lines are gains computed by the DGN and MLP, respectively. For both networks, gains were inferred almost perfectly after 15-20 minutes. B. Performance, measured as mean squared error between the true angular velocity, *Gs*(*t*) (Eq. (7)), and the angular velocity inferred by the networks. Same color code as panel A. C. Comparison of target angular velocity versus time (black) to that predicted by the DGN (blue). (A plot for the MLP is similar.) Before the gain change, the two are almost identical; immediately after the gain change, the network uses the previous gain. D. Top panel: Parallel fiber weights for the DGN network versus delay, *τ_i_* (Eq. (8)). Each panel shows 10 branches; 5 Purkinje cells are shown (chosen randomly out of 20). The weights vary smoothly with delay. Bottom panel: MLP weight profile, except that dendritic branches are replaced by the whole neuron (all 100 parallel fibers). The weights again vary smoothly with delay, but their shapes are now highly stereotyped.

Performance for both the DGN and the MLP were comparable and, after suitably adjusting the learning rates, the networks were able to learn in 15-20 minutes (Fig. 7A, B), consistent with learning times in behavioral experiments [79,80]. Figure 7C shows the target and predicted head velocities immediately before and after each gain change. Not surprisingly, immediately after a gain change, the network produces output with the old gain.

Figure 7D shows the connection strengths between parallel fibers (*x_i_*(*t*), Eq. (8)) and Purkinje cells, after learning, as a function of the delay, *τ_i_*. There are two notable features to these plots. First, the connectivity patterns are smooth. Second, although the DGN and the MLP solve the task equally well, there is a clear difference: for the MLP the connectivity patterns are highly stereotyped, while for the DGN they are far less so.

The smooth connectivity patterns, which are seen in both MLPs and DGNs, arise primarily because weights mediating inputs with similar delays have similar updates during learning. But there is another, somewhat technical, reason: the weights were initialized to small values. That’s important because for most directions in weight space, changes in the weights have no effect on the loss. Component of the weights that lie in these “null” directions will, therefore, not change with learning. Small initial weights ensure that the components in the null directions start small, and the lack of learning in these directions means they stay small.

The difference in the connectivity patterns – stereotyped versus diverse – are due to the fact that MLPs are not gated whereas DGNs are. The smooth, stereotyped connectivity patterns seen in MLPs arise because all neurons receive similar input statistics, and so they find similar solutions. The more diverse connectivity patterns seen in DGNs arise because inputs to different branches are gated differently, and so different branches do not see the same input statistics.

What happens when the initial weights are large and initially random? In that case, because the weights don’t change in the null directions, the final connectivity patterns are also not smooth – they’re almost as noisy as the initial weights. Here as well, though, there are difference between DGNs and MLPs: for DGNs the noise rides on top of diverse connectivity patterns very similar to those in the top panels of Fig. 7D, while for MLPs the noise is unmodulated, and more or less white (see Supplementary Fig. S4).

### 2.3 Testing predictions of the DGN in behaving animals

A key feature of DGNs is that the gates should exert a local effect on the dendritic branches of the principal neurons. When mapped onto the cerebellum, this suggests that the molecular layer interneurons (MLIs) should inhibit the individual dendritic branches of the Purkinje cells. We therefore tested whether individual Purkinje cell dendrites can be inhibited by activity in MLIs. Previous *in vitro* work has demonstrated that synaptic inhibition can locally inhibit calcium signals in Purkinje cell dendrites [46], and *in vivo* work has shown that MLI activity can influence the variability of these signals [48,61], but it has not yet been shown whether such effects can be localized to individual dendrites of Purkinje cells, and if so, what the spatial relationship of this effect is with presynaptic interneurons. Using multi-plane 2-photon imaging in awake PV-cre mice injected with the GCaMP7f virus, we could reliably record calcium signals from individual MLI somata and the climbing fiber-evoked calcium signals in the dendritic tree of Purkinje cells (Fig. 8A-B). With this approach, we were able to identify a substantial proportion of interneurons (72/142 MLIs in 3 mice; 51%) whose activation was associated with a significant decrease in climbing-fiber driven calcium signals in the dendrites of at least one nearby Purkinje cell(Fig. S5A). Given the axonal spread of MLIs [84,85], the analysis was confined to single interneurons that were located within 150 um (rostrocaudally) and 50 um (mediolaterally) from a given Purkinje cell dendrite. The extent of suppression varied between PC dendrites and also within individual PC dendrites recorded at different depths (Fig. 8C). In modulated Purkinje cell dendrites, the degree of suppression was 17.4 ± 0.5% (n = 133 Purkinje cells in 3 mice [range 6.6 to 53.5%]).

**Figure 8:**
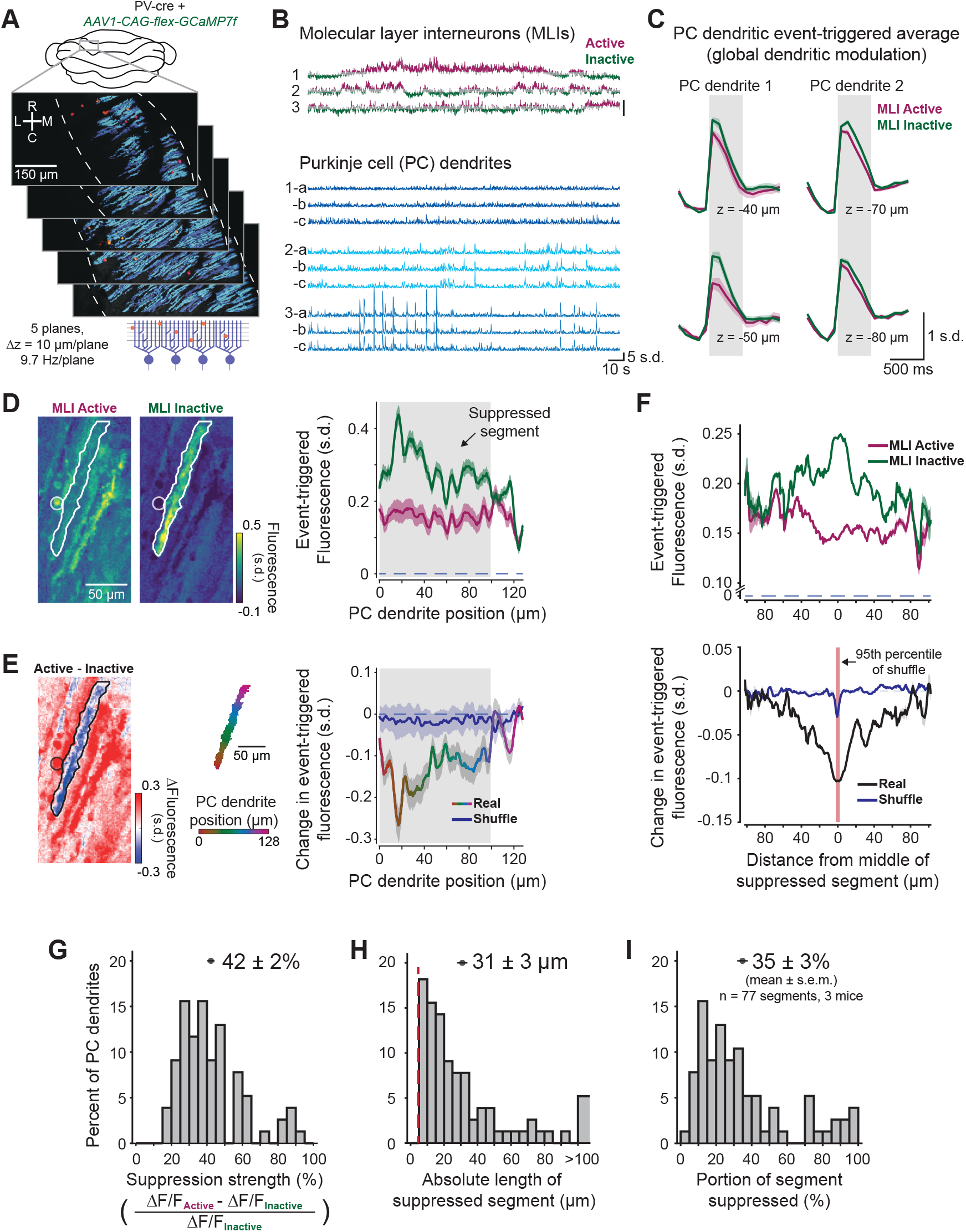
Dendritic gating of Purkinje cells by molecular layer interneurons *in vivo*. A. Multi-plane 2-photon calcium imaging of molecular layer interneurons (red) and Purkinje cell dendrites (blue). B. Example traces of MLIs (top, active and inactive states in purple and green) and Purkinje cell dendrites recorded in multiple planes marked a, b, and c (bottom). C. Two examples of mean climbing fiber-evoked Ca^2+^ signal, in two planes, in MLI-active (purple) and inactive (green) states. D. Spatial event-triggered map of area surrounding Purkinje cell dendrites (contoured region of interest) when a nearby MLI (circle projected from different plane) was in an active state or inactive state (left). Spatial profile of event-triggered fluorescence is shown on the right. E. Difference heatmap image (left) and spatial profile (rainbow trace, right) of event-triggered fluorescence of same dendrite shown in panel D. Shuffled trace (blue) was computed on a split of odd and even event indices. F. Mean spatial profile of event-triggered fluorescence in MLI-active and MLI-inactive condtions (top) and difference traces (bottom) aligned to middle of suppressed segment (black trace, *n* = 77 regions from *N* = 3 recordings in 3 mice). Shuffled trace (blue) was computed, as above, on an odd-even split. Suppressed regions smaller than the 95th percentile of the shuffle data (red region, 4.6 μm) were excluded. G. Histogram of strength of suppression calculated only within modulated segments. H. Histogram of suppressed segment lengths (red line shows 95th percentile of the shuffle). I. Histogram of suppressed segment lengths expressed as a percent of total segment length. Data are shown as mean ± S.E.M.

After identifying Purkinje cells whose global dendritic signals were inhibited when nearby MLIs were active, we investigated the spatial extent of this inhibition within Purkinje cell dendritic segments. We generated climbing fiber-evoked calcium signal maps in Purkinje cell dendrites, both when MLIs were active and when they were inactive (Fig. 8D, left). The difference revealed that MLI activation was associated with local suppression of calcium signals in subregions of the dendrites (blue region in Fig. 8E, left; Fig. S5B-E). To quantify the spatial extent of the suppression, we subdivided Purkinje cell dendritic regions into 1 *μ*m segments to generate spatial activity profiles in MLI-active and MLI-inactive conditions (Figs. 8D, right). Subtracting these yielded a spatial difference trace (Figs. 8E, right). Aligning these segments across all modulated PC dendrites allowed us to determine the average spatial profile of suppressed dendritic segments (Figs. 8F). To identify false positives, for each Purkinje cell dendrite analyzed we generated a shuffled difference trace, where MLI-active and MLI-inactive traces were replaced with an equal mixture of the two conditions (odd-even event split). We then identified suppressed regions in these shuffled difference traces, and used the 95th percentile (4.6 *μ*m) as the minimum length for a segment to be considered significantly modulated. Across all experiments, we identified *n* = 77 significantly modulated segments (*N* = 3 mice) whose calcium signals were suppressed by 42 ± 2% (mean ± S.E.M.) (Fig. 8G). The spatial extent of MLI-gated inhibition in these dendritic segments was 31 ± 3 μm (mean ± S.E.M.), accounting for 35 ± 3% (mean ± S.E.M.) of the extent of the dendritic tree at the imaged plane (Fig. 8H-I). These results demonstrate that MLIs can locally inhibit climbing fiber signals in the Purkinje cell dendritic tree with a spatial extent that is comparable to the length of individual spiny branchlets. This provides strong experimental evidence for a specific biological implementation of one of the principal design features of the DGN, namely local dendritic suppression of principal units by interneurons.

## 3 Discussion

Identifying the biologically plausible learning rules that mediate the modification of connections in neural networks is a key goal of both experimental and theoretical neuroscience. Here, we describe a new class of learning rules called Dendritic Gated Networks (DGNs). Each unit in each layer of a DGN consists of dendritic branches that are gated on and off by interneurons, and all units in all layers receive the same feedback learning signal. We show that the DGN has key advantages over existing learning algorithms, particularly in terms of learning speed and resilience to forgetting, tested across a range of learning tasks. We also show using *in vivo* experiments that key elements of the DGN architecture may be implemented in biological networks. These results suggest that DGNs may be widely useful in the machine learning community, and also suggest that this learning rule may be implemented in biological networks such as the cerebellum and other neural circuits with a similar network architecture.

### 3.1 Comparison of DGNs to other learning algorithms

The DGN network architecture differs from traditional learning algorithms (e.g. back-prop) as well as the algorithm on which it was based (Gated Linear Networks) in several important ways. Traditional learning algorithms like backprop map input to output in stages, with the input gradually transformed, until eventually, in the output layer, the relevant features are easy to extract. There is certainly some evidence for this hierarchical strategy being implemented in the brain. It is, for example, much harder to extract which face a person is looking at from activity in visual area V1 than in fusiform face area [86, 87]. While this strategy for computing is reasonable, it has a downside: the relationship between activity in intermediate layers and activity in the output layer is highly nontrivial, which makes it especially hard for the brain to determine how weights in intermediate layers should change.

Despite the inherent complexity of this strategy, biologically plausible learning rules that implement it have been proposed [15–27]. The DGN algorithm takes a different approach from any of these. With this architecture, dendritic branches are gated on and off via a random (and fixed) linear transformation of their input. The summed activity of these branches forms the prediction of the neuron, which gets adjusted over time via a delta rule. Consequently, all neurons predict the same target; and each layer improves upon the predictions of the previous layer.

The DGN also differs from and improves over the algorithm on which it was based – the Gated Linear Network (GLN) [29, 30]. In particular, the GLN requires a bank of weights for each neuron, with the input choosing which one the neuron should use – something that seems extremely difficult for the brain to implement. The DGN, however, replaces the library of weights with gated dendritic branches, an innovation essential for biological plausibility. Thus, although the DGN is conceptually related to the GLN, from the point of view of neuroscience it has a critical new component which makes it, unlike the GLN, relevant to the brain.

### 3.2 Implementation in cerebellar circuits

The architecture and exceptional efficiency of learning exhibited by DGNs suggests that this algorithm may also be implemented in biological networks. Specifically, several key features of the DGN are recapitulated in the functional architecture of the cerebellum. First, the cerebellum receives a clear and global feedback signal in the form of the climbing fiber input to Purkinje cells and cerebellar nuclear neurons that is the principal driver of learning in the cerebellar circuit [88]. Second, the principal neurons of the cerebellar cortex, Purkinje cells, exhibit linear encoding of their inputs due to their high baseline firing rates and unique biophysical properties [36, 89]. Finally, molecular layer interneurons, which are known to target dendritic branches of Purkinje cells, are likely candidates to mediate branch-specific dendritic inhibition [46, 48, 90].

A key prediction of the DGN that would bolster its biological plausibility is that interneurons should gate activity in single dendritic branches of principal cells. Here, we provide the first *in vivo* evidence that molecular layer interneurons can produce inhibition of dendritic calcium signals on the level of single dendritic branches in Purkinje cells, a longstanding, but until now untested, prediction of anatomical [91–94] and theoretical [63, 90, 95] work. By simultaneously imaging dendritic calcium signals in Purkinje cells and activity in neighboring MLIs, we show that MLI activity can substantially decrease dendritic calcium signals in Purkinje cells. Previous *in vitro* work has shown that even modest inhibition of dendritic calcium signals (on order of 20%) can completely abolish cerebellar plasticity [96], suggesting that the suppression we observe is capable of abolishing learning. These MLI-driven effects were not distributed equally across PC dendritic arbors. Indeed, suppression of dendritic calcium signals was often restricted to individual dendritic branches of Purkinje cells, even as neighboring regions of the same dendritic arbor were unaffected. The profound local suppression of these climbing fiber-driven signals suggest that MLI-driven inhibtion is also capable of suppressing parallel fiber-driven input to Pukrinje cells, which are relatively much weaker [97]. Thus, it is likely that MLIs can gate both input and learning to single Purkinje cell dendritic branches. In summary, our demonstration that MLIs can modulate the Purkinje cell dendritic calcium signals on the spatial scale of a single dendritic spiny branchlet strongly supports the DGN gating prediction. We note that the binary on/off gating exhibited by the DGN is challenging to implement biologically; far more likely are softer gates, where the amount of gating is a function of the input. To determine how soft gates affected DGN performance, we performed simulations on the permuted MNIST and inverse kinematics tasks, and the results were virtually identical to the ones with on/off gates (data not shown), emphasizing the flexibility of DGN implementation in the brain. Our experimental data (together with previous work linking cerebellar functional architecture to features of the DGN) provides strong support for the idea that a DGN-like algorithm is implemented in cerebellar circuits.

The architecture of the DGN makes several further predictions about how the DGN may map onto the cerebellum. A key prediction that awaits experimental validation is that the activity of MLIs should depend predominantly on parallel fiber input and the input-output relationship of these interneurons should change very slowly relative to the timescale over which Purkinje cells learn, which can be measured in single trials [98]. Assessing the stability of the parallel fiber-mediated input-output relationship of MLIs is difficult because granule cells are known to exhibit learning-related changes in activation [99,100]. Thus, further experiments to determine the stability of parallel fiber-MLI and parallel fiber-Purkinje simple spike input-output relationships over learning will need to account for changes in the firing patterns of cerebellar afferents. Another prediction of the DGN is that the parallel fiber connectivity pattern in the VOR or similar tasks should carry information about architecture. If parallel fiber-Purkinje cell connectivity is smooth and diverse (Fig. 7D, top panels), this supports a DGN-like implementation. If, on the other hand, connectivity is smooth and stereotyped (Fig. 7D, bottom panels) or noisy with no other temporal structure (Fig. S4), this argues for alternative implementations, such as an MLP. Distinguishing between these alternatives experimentally would require recording from populations of individual granule cells during VOR learning, then mapping the synaptic strength between those same granule cells and Purkinje cells.

### 3.3 DGNs in other neural circuits

While DGNs exhibit features that make them particularly well-suited for implementation in the cerebellum, the general principles of this learning rule may be applicable to a variety of brain circuits. In particular, the gating of dendritic signals that we demonstrate in Purkinje cells may also be a feature of cortical networks, given the branch-specific innervation of interneuron axons that has been documented in many cortical circuits [101]. Such generalization would require some modifications to implement the algorithm; for instance, the learning rule will have to change because activity in the cortex is far from linear.

### 3.4 Conclusions

In summary, Dendritic Gated Networks are strong candidates for biological networks – and not just in the cerebellum; they could be used anywhere there is approximately feedforward structure. They come with two highly desirable features: rapid, data-efficient learning, and biologically plausible learning. Furthermore, they suggest a novel role for inhibitory neurons, which is that they are used for gating dendritic branches. We anticipate that the strong, experimentally testable, predictions may inspire investigations in many brain circuits where rapid learning may invoke a DGN algorithm.

## 4 Methods

### 4.1 Classification tasks

The network we use in our model is described in Eqs. (1) and (3), and the learning rules are given in Eq. (4). For regression (VOR and inverse kinematics), we use the identity function for *ϕ* (Eq. (1)), and the square loss (Eq. (5)), resulting in the update rule given in Eq. (6). For classification (the simple task described in Fig. 3 and permuted MNIST), the network computes probabilities, so *ϕ* needs to be bounded, and a square loss is not appropriate. Here we provide details for this case.

For classification we use a standard sigmoid function, *σ*(*z*) = *e^z^*/(1 + *e^z^*), albeit modified slightly,

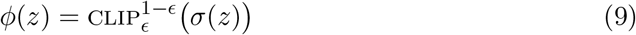

where 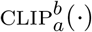 clips values between *a* and *b* (so the right hand side is zero if *σ*(*z*) is smaller than e or larger than 1 – *ϵ*). Clipping is used for bounding the loss as well as the gradients, which helps with numerical stability. It also enables a worst-case regret analysis [29,30]. We set *ϵ* to 0.01, so neural activity lies between 0.01 and 0.99.

The square loss is not appropriate in this case, so instead we use the binary cross-entropy loss: the loss of neuron *i* in layer *k* is given by

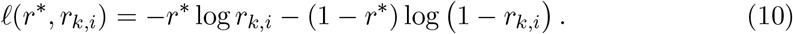

Consequently, the update rule for the weights, Eq. (4), is (after a small amount of algebra)

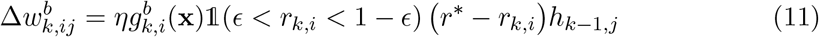

where 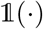 is 1 when its argument is true and 0 otherwise. The fact that learning is zero when *r_k,i_* is outside the range [*ϵ*, 1 – *ϵ*] follows because *dϕ*(*z*)/*dz* = 0 when *z* is outside this range (see Eq. (9)). This ensures that learning saturates when weights become too large (either positive or negative). However, this can cause problems if the output is very wrong; that is, when *r*\* = 1 and *r_k,i_* < *ϵ* or *r*\* = 0 and *r_k,i_* > 1 – *ϵ*. To address this, we allow learning in this regime. We can do that compactly by changing the learning rule to

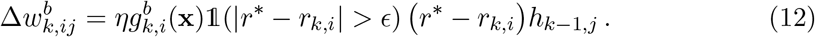

Essentially, this rule says: stop learning when *r_k,i_* is within *ϵ* of *r*\*.

For a compact summary of the equations (given as pseudocode), see Supplementary Algorithms S1 and S2.

### 4.2 Simulations

Here we provide details of our simulations. Simulations were written using JAX [102], the DeepMind JAX Ecosystem [103], and Colab [104].

#### Permuted MNIST

We adopt the pixel-permuted MNIST benchmark [57, 60], which is a sequence of MNIST digit classification tasks with different pixel permutations. Each task consists of 60,000 training images and 10,000 test images; all images are deskewed. Models are trained sequentially across 10 tasks, performing a single pass over all 60,000 training examples for each of the tasks. We provide the implementation details below; the parameters swept during a grid search are given in Supplementary Table S2.

##### DGN

We use networks composed of 100 and 20 units in the hidden layers and a single linear neuron for the output. All neurons in the hidden layers have 10 dendritic branches. The output of the network is determined by the last neuron. MNIST has 10 classes, each corresponding to a digit. Therefore, we use 10 DGN networks, each encoding the probability of a distinct class. These networks are updated during training using a learning rate *η* = 10^-2^. During testing, the class with the maximum probability is chosen. Images are scaled and shifted so that the input range is [–1, 1]. The gating vectors, 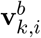, are chosen randomly on the unit sphere, which can be achieved by sampling from an isotropic Normal distribution and then dividing by the L2 norm. The biases, 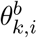, are drawn independently from a zero mean Gaussian with standard deviation 0.05.

##### MLP and EWC

We use a ReLu network with 1000 and 200 neurons in the hidden layers and 10 linear output units with cross entropy loss. In this setting, the MLP and EWC have the same number of neurons as the DGN, but fewer connections. We use the ADAM optimization method [105] with a learning rate *η* = 10^-4^ (see Supplementary Table S2 for details of the hyperparameter optimization), in conjunction with dropout. We use mini-batches of 20 data points. For EWC, we draw 100 samples for computing the Fisher matrix diagonals and set the regularization constant to 10^3^.

#### Inverse Kinematics

Each DGN network has 20 Purkinje cells and one linear, non-gated, output neuron, and we vary the number of branches. We use a quadratic loss, as in Eq. (5), a learning rate *η* = 10^-5^, and we run for 2000 epochs (2000 passes over the dataset). The inputs are centered at 0 and scaled to unit variance per dimension, and the targets are scaled so that they lie between 0 and 1. The reported MSEs are computed on the test set based on inverse transformed predictions (thus undoing the target scaling). The gating parameters are chosen in the same way as for the MNIST simulations described above.

We discovered that the training set of the SARCOS dataset (downloaded from http://www.gaussianprocess.org/gpml/data/ on 15 December 2020) includes test instances. To the best of our knowledge, other recent studies using the SARCOS dataset [106, 107] reported results with this train/test setting. This means that the reported errors are measures of capacity rather than generalization. We compare the performance of DGN against the best known SARCOS results in Supplementary Table S1 using the existing train/test split.

#### VOR

The gating parameters 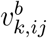 and 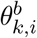 (Eq. (3)), were drawn independently from the standard normal distribution. The learning rates were *η* = 10^-5^ for the DGN and *η* = 0.03 for the MLP.

### 4.3 Animal experiments

#### Animal housing and surgery

All animal procedures were approved by the local Animal Welfare and Ethical Review Board and performed under license from the UK Home Office in accordance with the Animals (Scientific Procedures) Act 1986 and generally followed procedures described previously [108]. Briefly, we used PV-Cre mice (B6;129P2-Pvalbtm1(cre)Arbr/J) [109] maintained on a C57/BL6 background. Mice were group housed before and after surgery and maintained on a 12:12 day-night cycle. Surgical procedures were similar to those described in [108], except that we injected Cre-dependent GCaMP7f (pGP-AAV-CAG-FLEX-jGCaMP7f-WPRE [serotype 1]; [110]) diluted from its stock titer to a final concentration of 3 x 10^11^ GC/ml (~1:25). After mice had recovered from surgery, they were acclimated to the recording setup and expression-checked before beginning recordings.

#### Two-photon calcium imaging data acquisition and processing

Imaging experiments were performed using a 16x/0.8 NA objective (Nikon) mounted on a Sutter MOM microscope equipped with the Resonant Scan Box module. A Ti:Sapphire laser tuned to 930 nm (Mai Tai, Spectra Physics) was raster scanned using a resonant scanning galvanometer (8 kHz, Cambridge Technologies) and images were collected at 512×256 pixel resolution over fields of view of 670×335 *μ*m per plane at an average power of 30-70 mW. Volumetric imaging across 5 planes spaced by 10 *μ*m (40 μm depth range per recording, total depth ranging 10-70 *μ*m below pial surface) were performed using a P-726 PIFOC High-Load Objective Scanner (Physik Instruments) at an effective volume rate of 9.7 Hz. The microscope was controlled using ScanImage (Version 5.6, Vidrio Technologies) and tilted to 12.5 degrees such that the objective was orthogonal to the surface of the brain and coverglass.

Regions of interest (ROIs) corresponding to single MLI somata and Purkinje cell dendrites, which were easily distinguishable based on their shape, were identified using Suite2p software [111] for initial source extraction and custom-written software for subsequent analyses. Fluorescence time series were computed as (*F* – *F*0)/*F*0 where *F* was the signal measured at each point in time and F0 is the 8th percentile of a rolling average baseline surrounding each data point (2000 frames for MLIs and 10 frames for Purkinje cell dendrites). A neuropil correction coefficient of 0.5 (50 percent of neuropil signal output from Suite2p) was applied to MLI ROIs. A range of baseline durations and neuropil correction coefficients were tested and varying these parameters did not alter the main findings. Following these calculations, fluorescence signals were *z*-scored to facilitate comparisons across neurons. Signals from the same Purkinje cell dendrites recorded in multiple planes were identified based on a correlation threshold (*>*0.5, followed by manual confirmation) and analyzed independently for dendritic modulation experiments. Event times in dendrites were detected using MLspike [112].

#### MLI gating of Purkinje cell dendritic signals

For dendritic modulation experiments, active and inactive MLI states were defined as imaging frames where activity in an MLI deviated more than 0.5 standard deviations above or below the mean, respectively (Fig. S5). Using a higher threshold yielded similar results but resulted in fewer dendritic events in each condition. After identifying these states, we compared the magnitudes of isolated dendritic events in these two conditions for Purkinje cells within a 300 × 100 μm ellipse centered on each MLI whose major axis is parallel to that of Purkinje cell dendrites in the field of view (approximately rostrocaudal). Ellipse dimensions were chosen to approximate the known rostro-caudal and mediolateral spread of MLI axons [84, 85]. Only isolated events in Purkinje cell dendrites (defined as those that occurred more than 500 ms before and after any other events) were analyzed, and fluorescence event magnitudes were calculated over the 5 frames (~500 ms) after each event for initial identification of MLI-modulated dendrites. Because analysis of each recording involved many thousands of comparisons, we assessed significance differences between Purkinje cell dendrite event magnitudes in MLI active and inactive states with a significance threshold of 0.05 that was corrected for multiple comparisons using false discovery rate threshold of 5% [113].

Motion-corrected fluorescence movies that were used for pixel-wise analysis of dendritic subregions were pre-processed by first correcting for slow fluctuations in fluorescence, which was done by *c*omputing (*F* – *F*_0_)/*F*_0_ where *F* was the signal measured at each point in time and *F*_0_ is the 8 percentile of a rolling average baseline surrounding each data point (2000 frames), and then *z*-scoring. Purkinje cell dendritic ROIs defined in Suite2p were segmented into 1 *μ*m increments by fitting a 4th degree polynomial to each ROI, grouping ROI pixels closest to regular spaced points along this fit line, computing a weighted average of these pixels based on the pixel weights assigned by Suite2p, and smoothing over 5 *μ*m. Activity profiles in MLI active and inactive states were subtracted and used to generate spatial suppression profiles for each Purkinje cell dendrite and error bars were generated from the summed variances in active and inactive conditions. Shuffled distributions were generated by replacing active and inactive conditions with an odd-even event split of each of these groups, yielding two distributions comprising of 1/2 MLI-active and 1/2 MLI-inactive events for each dendritic segment. Significantly modulated dendritic segments were defined as the longest region for each dendritic segment in which the 95% confidence interval of the difference trace was less than zero. To account for false positive dendritic segments that would be identified by finding minima in noise, we performed this identification procedure on our shuffled data and excluded identified dendritic segments in our real data that were smaller than the 95th percentile of these fictive segments (4.6 *μ*m).

## Code availability

We provide pseudo code in Supplementary Algorithms S1 andS2. A simple python implementation can be accessed via https://github.com/deepmind/deepmind-research/blob/master/gated_linear_networks/colabs/dendritic_gated_network.ipynb.

## Data availability

The data that support the findings of this study are available from the corresponding authors upon reasonable request. Additional analysis made use of standard publicly available benchmarks including MNIST [114] and SARCOS (http://www.gaussianprocess.org/gpml/data/).

## Acknowledgements

We thank Timothy Lillicrap, Gregory Wayne, Eszter Vértes, and Brendan Bicknell for valuable feedback. Michael Häusser is supported by the Wellcome Trust and the European Research Council. Peter Latham is supported by the Gatsby Charitable Foundation and the Wellcome Trust.

## Author contributions

ES and AGB developed the computational model with advice from JV, CC, PEL, DB, and MHut. AGB, ES, and SK performed simulation experiments and analysis with advice from CC and PEL. DK and MBea acquired and analyzed neuronal data with advice from MHäu. PEL, AGB, ES, and DK wrote the paper with help from all other authors. AGB and ES managed the project with support from MBot, CC, JV, and DB.

## Supplementary Information

### Difference between GLNs and DGNs

The main difference between the DGNs and Gated Linear Networks (GLNs) is in the interplay of the gating functions and synaptic weights. To understand this, it is helpful to write Eq. (1) in the form

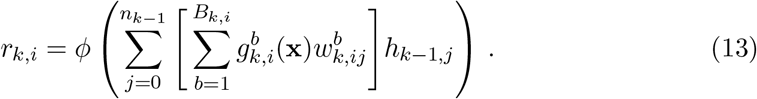

The term in brackets is the effective weight, which, for each *j*, consists of a sum over branches. The gates, 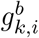, can be either zero or 1; since there are *B_k,i_* of them, there are 2^*B_k,i_*^ possible effective weights. For the GLN, on the other hand, *r_k,i_* is given by

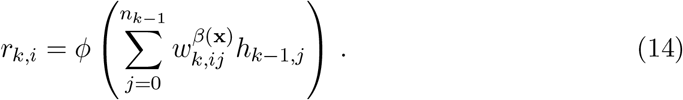

The difference between this and Eq. (13) is the the term in brackets has been replaced by a single weight, 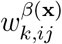. However, the index *β*(**x**) can take on 2^*B_k,i_*^ values, so there are just as many weights in the GLN as effective weights in the DGN. The value of *β*(**x**) is determined by the input, and is given by the binary string (suppressing the **x** dependence for clarity)

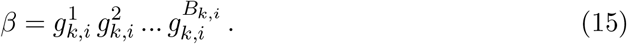

(So if there were 5 gates, and the 3rd and 5th were on while the others were off, the term on the right would be 00101, and *β* would be 5.)

In our experience, for similar B, DGNs and GLNs perform equally well. Computationally, DGNs are more memory efficient, as B weight vectors need to be stored per neuron as opposed to 2^*B*^ for GLNs. However, this comes at the cost of more operations, as there is an additional sum over branches (the term in brackets in Eq. (13)).

The difference in the number of parameters translates to a difference in inductive bias. GLNs are less prone to catastrophic forgetting compared to DGNs, as only one weight vector per neuron is updated for each input. This, however, means that DGNs are better than GLNs at learning new tasks – so long as there is some shared structure.

### Convexity

Here we show that the loss is convex with respect to the weights in the previous layer. Temporarily dropping indices for clarity, the loss, *ℓ*(*r*,r*), is given in terms of the weight vector, **w**, as *ℓ*(*r*,r*) = *ℓ*(*r*\*, *ϕ*(*h*)) with *h* = **c** · **w** (see Eq. (1)). If *ℓ*(*r*\*, *ϕ*(*h*)) is convex in *h*, then *ℓ* is also convex in **w**, since *h* is a linear function of **w**.

For quadratic loss, Eq. (5), *ϕ* is the identity, so 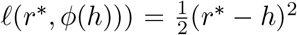. This is obviously convex in *h*, and so convex in **w**. For binary cross-entropy loss, Eq. (10), *ϕ*(*h*) is given by a clipped sigmoid, Eq. (9). When clipped, *ϕ*(*h*) = 0, which is convex. When not clipped, *ϕ*(*h*) = *σ*(*h*) = 1/(1 + *e*^-*h*^), for which it is easy to show that *∂*^2^ℓ(*r*\*, *ϕ*(*h*)))/*∂h*^2^ = *σ*(*h*)(1 – *σ*(*h*)) > 0. Thus, again *ℓ* is convex in *h*, and so also in **w**.

### Optimal number of branches

The number of dendritic branches is one of the main factors determining the model capacity of DGNs. Having too few branches results in underfitting, as the network is not flexible enough to learn the underlying function. Having too many branches, on the other hand, can result in memorization, and thus overfitting. We have generally used 10 branches per neuron except in the Inverse Kinematics experiments, where we used up to 5000 branches, as the task measures memorization not generalization.

In Figure S1, we show the average accuracy in the permuted MNIST task as a function of the number of branches. This inverted U-shaped relationship can be observed in most tasks (data not shown). In Figure S2, we show how the MSE improves with an increased number of branches. Because the training and test data is mixed, over-fitting is not possible, and so performance improves monotonically with the number of branches.

**Figure S1:**
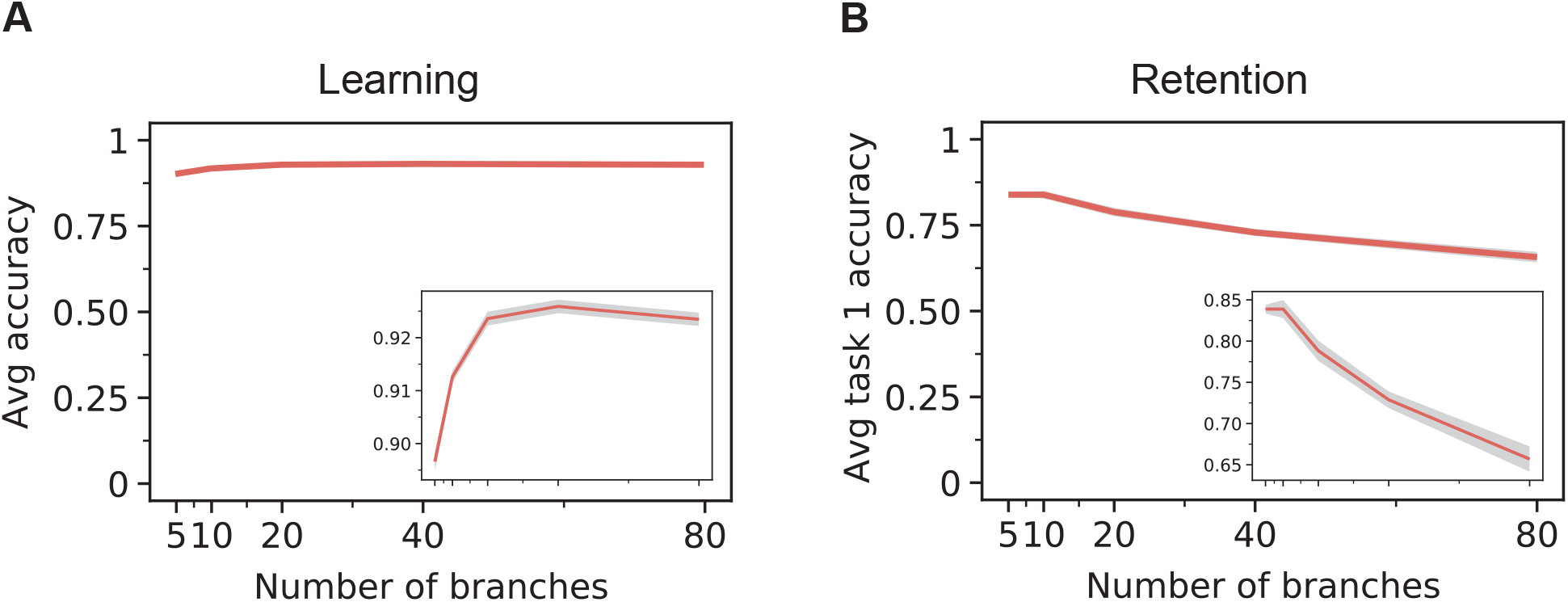
Permuted MNIST as a function of the number of DGN branches. A. Test accuracy at the end of training of each task, averaged over all 10 tasks. B. Test accuracy on task 1 after training on all 9 permutations. Grey areas are 99.5% confidence intervals of the results obtained from 10 models, initialised with different gating parameters and trained on differently permuted data.

**Figure S2:**
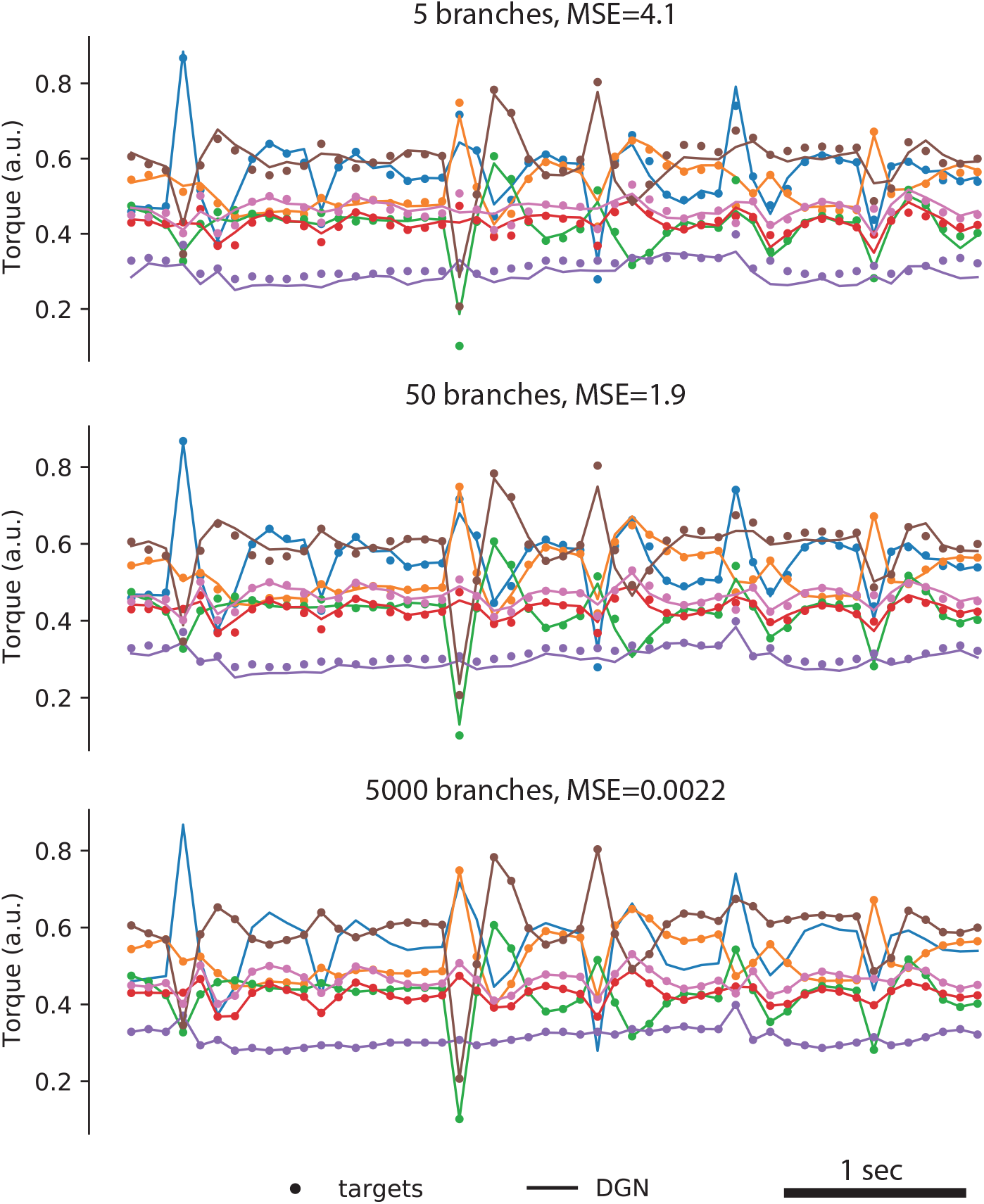
Sarcos solution for DGNs with 5, 50 and 5000 branches. Learning rate was 10 ^−4^ for 5 branches and 10^-5^ for 50 and 5,000 branches.

### Inverse Kinematics

In Table S1 we compare the mean square error (MSE) obtained by DGN against baselines obtained from [31,106,107]. Note that, as mentioned in Methods, we (like others) used a test set that contained training examples.

**Table S1:** Test mean square error (MSE) and the number of passes over the dataset (i.e., number of epochs) for DGN with 5,000 branches versus previously published methods on the SARCOS inverse dynamics dataset [76,106,107]. DGN obtains the best result, by a factor of 50.

| Algorithm | MSE | Epochs |
| --- | --- | --- |
| DGN | **0.002** | 2000 |
| Random forest | 2.39 | - |
| MLP | 2.13 | - |
| Stochastic decision tree | 2.11 | - |
| Gradient boosted tree | 1.44 | - |
| TabNet-S | 1.25 | 55000 |
| Adaptive neural tree | 1.23 | - |
| TabNet-M | 0.28 | 55000 |
| TabNet-L | 0.14 | 55000 |
| Gaussian Gated Linear Network | 0.10 | 2000 |

### Catastrophic Forgetting (permuted MNIST)

Hyerparameter selection. We used a grid search to select the hyperparameters for the three networks (DGN, MLP and EWC). The parameters we tested are shown in Table S2; the ones that maximize test accuracy are in bold.

**Table S2:**
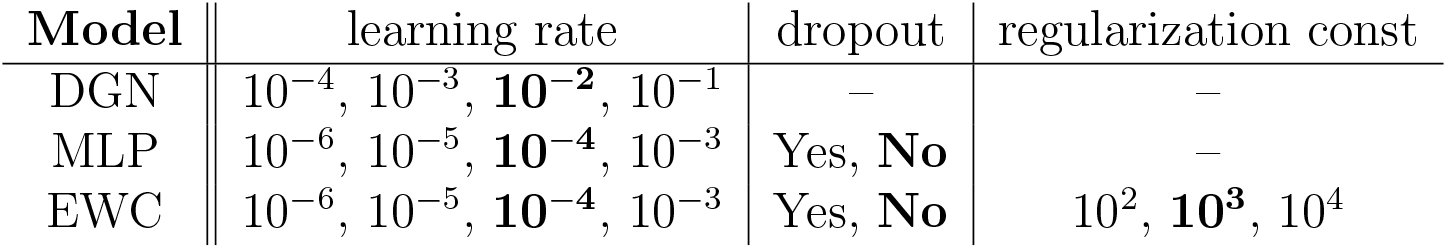
For permuted MNIST, parameters swept during grid search. The best parameters (shown in bold) are the ones that maximize the average test accuracy over 20 random seeds.

| Model | learning rate | dropout | regularization const |
| --- | --- | --- | --- |
| DGN | $10^{-4}$, $10^{-3}$, **$10^{-2}$**, $10^{-1}$ | – | – |
| MLP | $10^{-6}$, $10^{-5}$, **$10^{-4}$**, $10^{-3}$ | Yes, **No** | – |
| EWC | $10^{-6}$, $10^{-5}$, **$10^{-4}$**, $10^{-3}$ | Yes, **No** | $10^2$, **$10^3$**, $10^4$ |

#### Learning curves

In Fig. S3 we show the test performance of previously learned tasks (columns) as a function of the training across multiple tasks. To reduce clutter, a subset of the tasks (1, 2, 4, and 8, out of 10) are shown. The top left plot (train and test on task 1) shows that DGNs learn the first task much faster than all other methods. The plots to the right of that show retention on task 1 while the network is sequentially trained on subsequent tasks. MLP performances drop drastically after learning a few new tasks, while DGN and EWC show little forgetting. This is a remarkable feat for DGNs, which have no access to task boundaries and no explicit memory of previously learned tasks. EWCs, on the other hand, have both. If we look at the four diagonal plots, we see that DGN learns new tasks faster than all other methods, although the difference gets smaller as more tasks are learned.

The final accuracies across the diagonal correspond to the left panel of Fig. 4 whereas the final accuracies across the first row correspond to the right panel.

### VOR

To obtain the smooth connectivity patterns seen in Fig. 7D, the initial weights had to be small. Larger initial weights produced non-smooth connectivity patterns, although the non-smoothness was different for MLPs than it was for DGNs. For MLPs, standard Glorot intialisation led to the noisy connectivity patterns shown in Fig. S4D, bottom panel; in contrast, to produce smooth patterns, the weights had to be scaled down by a factor of of 100. For DGNs, scaling the initial weights up by a factor of 10 relative to Fig. 7D produced noisy weights, but riding on a smooth background (Fig. S4D, top).

**Figure S3:**
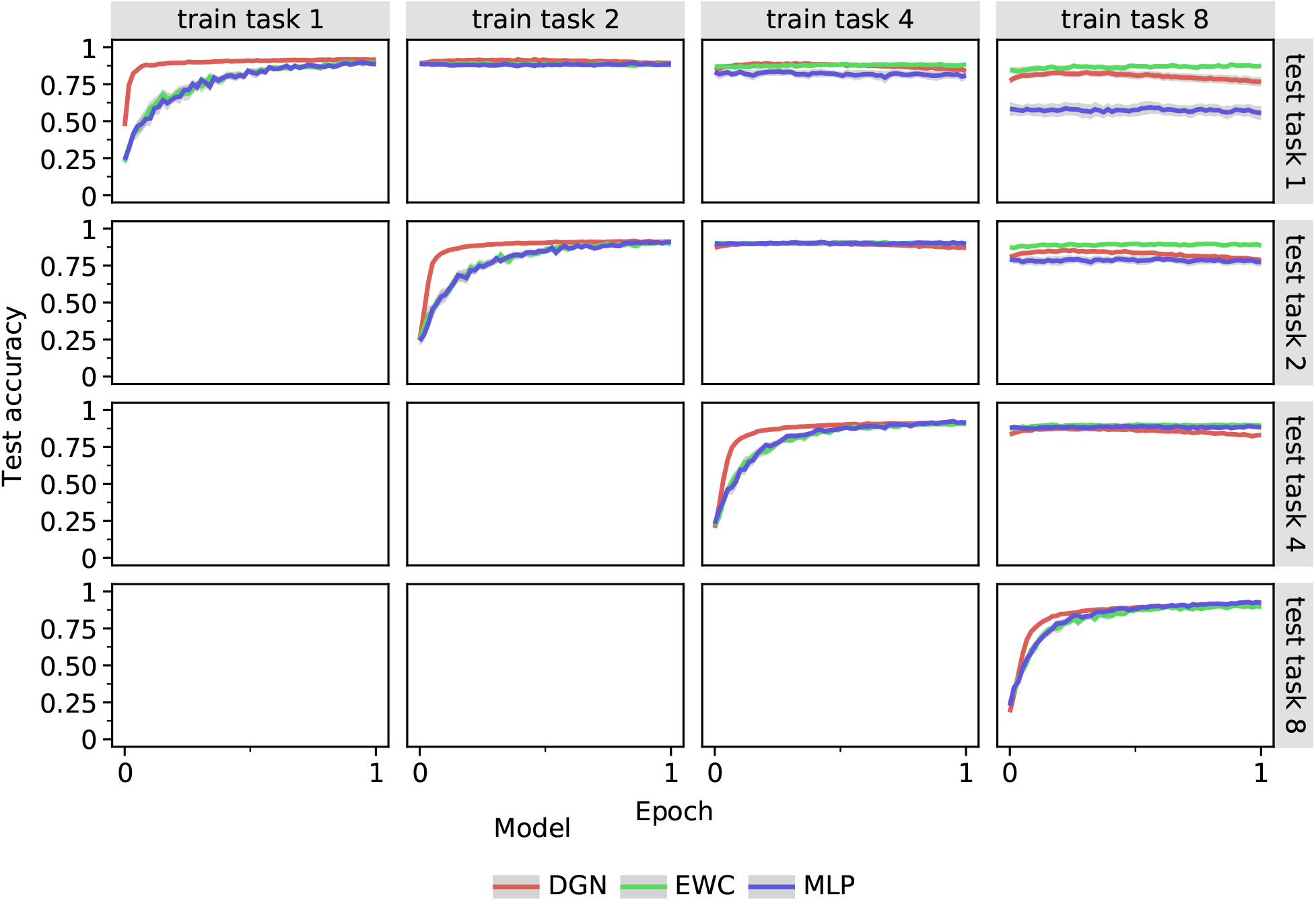
Retention results for permuted MNIST. Models are trained sequentially on 10 tasks, a subset of which is shown (tasks 1, 2, 4 and 8). Each column corresponds to a different stage of training (see labels on top), and each row reports test accuracy for a specific task. For example, the top row indicates performance on task 1 after being trained sequentially on tasks 1, 2, 4 and 8. Each model trains for one epoch per task; i.e., the 60,000 training examples per task are used only once. Error bars, indicated by the thickness of the lines, denote 95% confidence levels over 20 random seeds.

**Figure S4:**
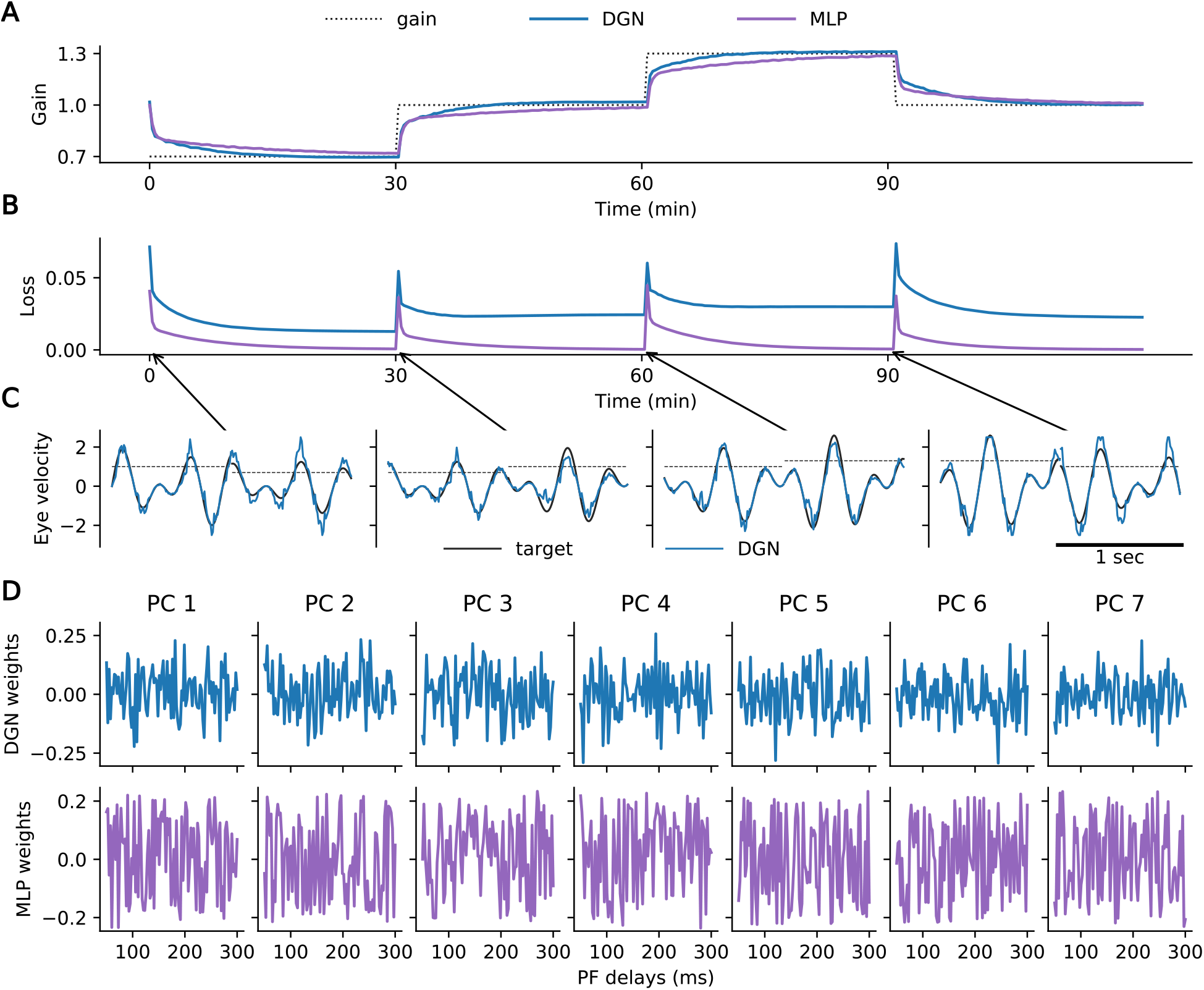
VOR network initialized with large, noisy weights. Same as Fig. 7, except that the training starts from large, noisy weights. For clarity, only five branches are shown in the top panel of D (compared to 10 in Fig. 7).

**Figure S5:**
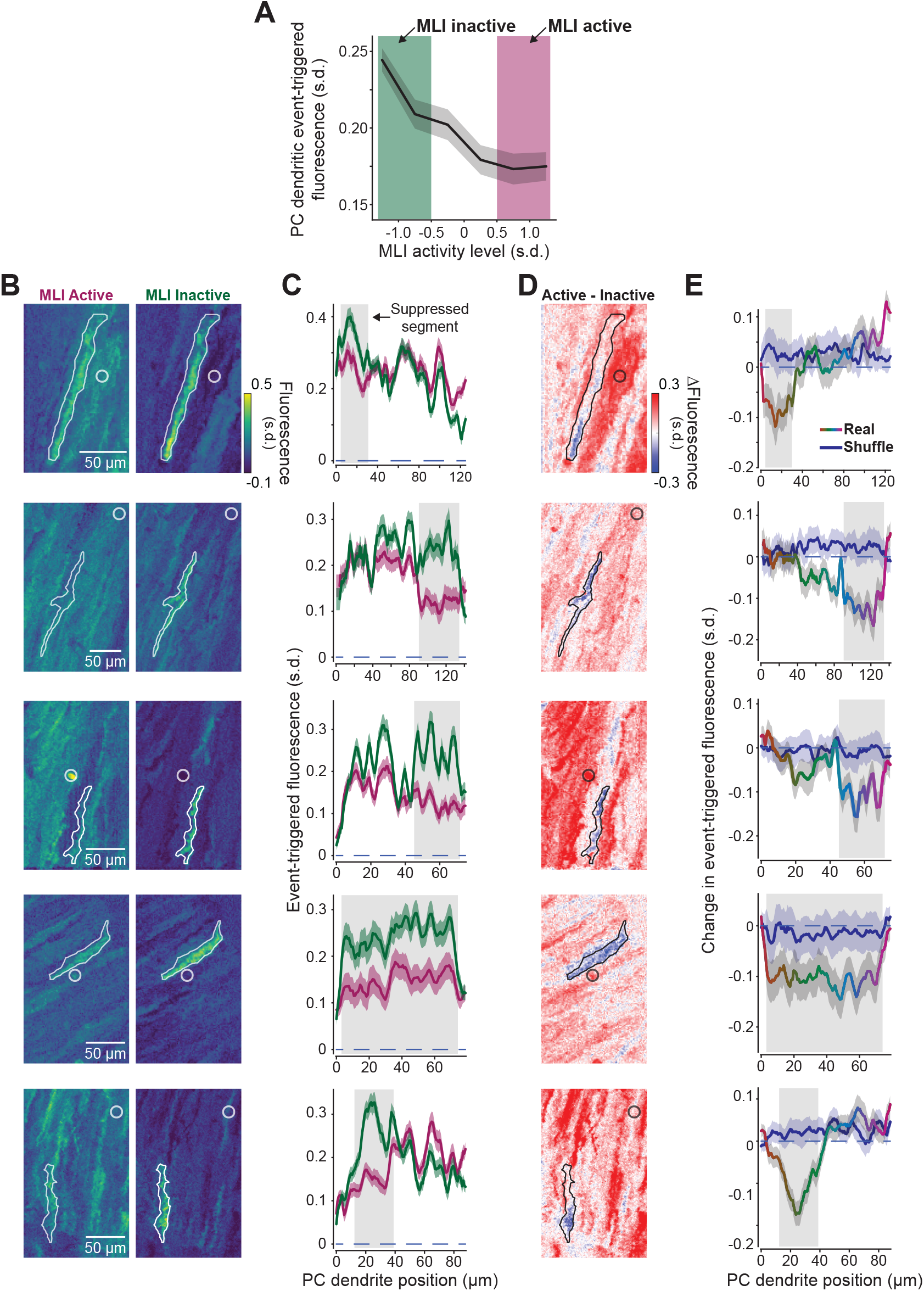
Suppression of Purkinje cell dendritic segments by MLIs. A. Event-triggered fluorescence in modulated Purkinje cell dendritic segments as a function of MLI activity level. Activity levels used for analysis in main figure are highlighted. B. Five additional examples of spatial event-triggered map of area surrounding Purkinje cell dendrites (contoured region of interest) when a nearby MLI (white circle; sometimes projected from different plane) was in an active state or inactive state. C. Spatial profile of event-triggered fluorescence of PC dendrites shown in panel A. D. Difference heatmap image of event-triggered fluorescence of same dendrites shown in panels A-B. E. Spatial profile trace (rainbow) and shuffled trace (blue) of of event-triggered difference heatmap images.

### Pseudocode

#### Algorithm S1 DGN for quadratic loss

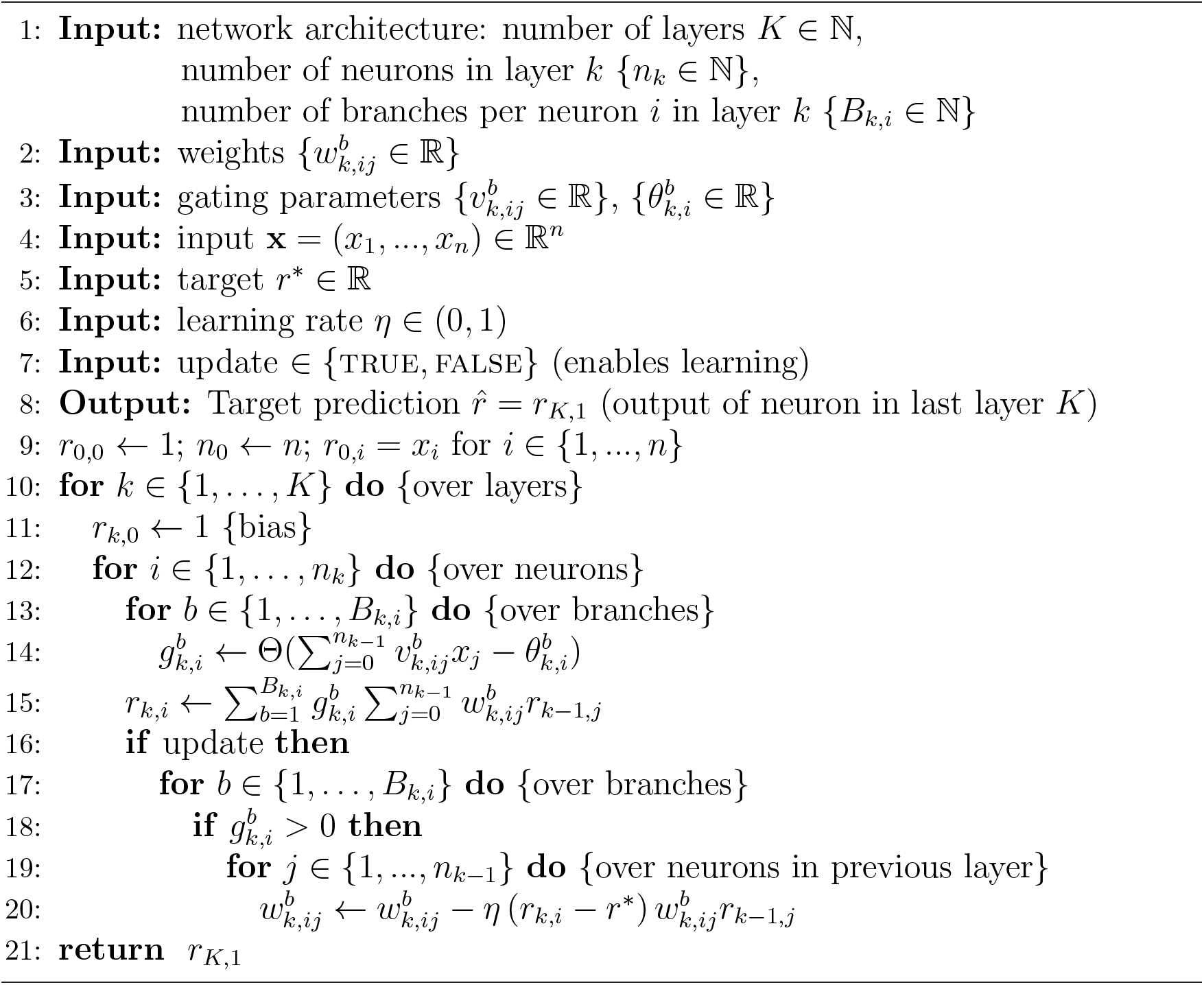

Here *Θ*(·) is the Heaviside step function (Θ(*z*) = 1 for *z* > 0 and Θ(*z*) = 0 otherwise).

#### Algorithm S2 DGN for Bernoulli data

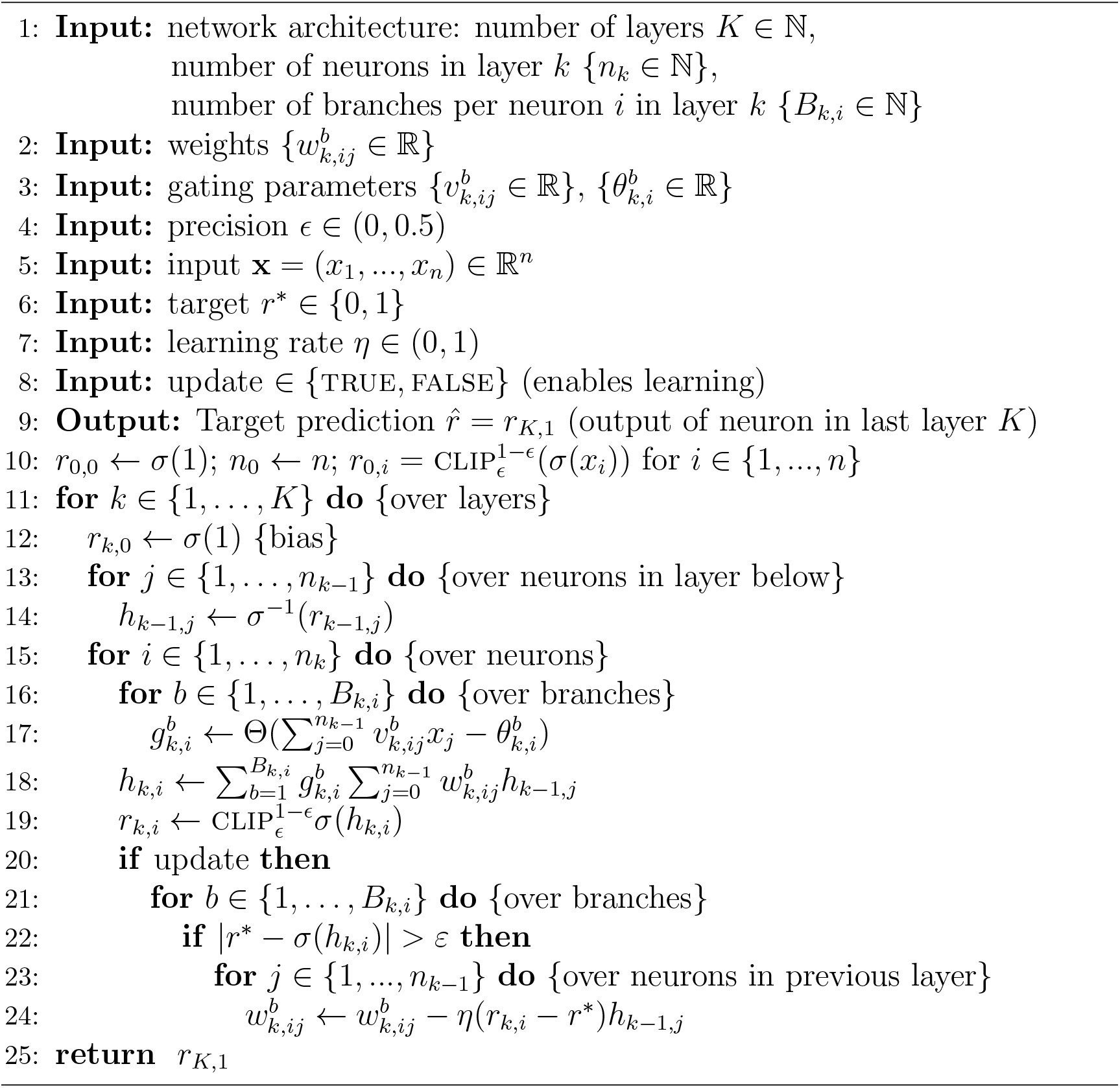

Here, as above, 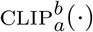 clips values between *a* and *b*,

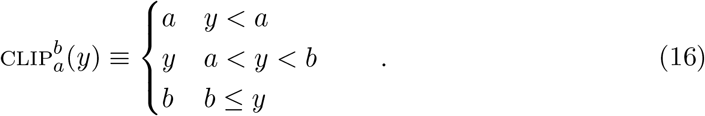

Also as above, *σ*(·) is the sigmoid function, *σ*(*z*) = *e^z^*/(1 + *e^z^*). Its inverse is given by *σ*−1(*y*) = log(*y*/(1 – *y*)).

